# CSF-Seq enables transcriptome-wide profiling of cerebrospinal fluid and identifies prognostic signature of leptomeningeal disease

**DOI:** 10.64898/2026.05.21.725787

**Authors:** Maxine Umeh Garcia, Giuseppe Barisano, Pablo Nunez Perez, Thy Trinh, Rukayat Taiwo, Daniel Herrick, Meaghen Roy-O’Reilly, Seunghyun Lee, Elias Spiliotopoulos, Charlotte Weixel, Georgiana Burnside, Bryana Godfrey, Yi Zhang, Sophia Chernikova, Sofia Tosoni, Monica Granucci, Cecile Riviere-Cazaux, Gwen Coffey, Eleanor Villanueva, Terry C. Burns, Seema Nagpal, Thuy T.M. Ngo, Melanie Hayden Gephart

## Abstract

Leptomeningeal disease (LMD) is a rapidly fatal complication of systemic cancer for which sensitive diagnostic tools and informative biomarkers remain limited. Here, we introduce CSF-Seq, a method for whole-transcriptome sequencing of cell-free RNA (cfRNA) from human cerebrospinal fluid (CSF), designed to enable molecular profiling of LMD and other central nervous system (CNS) conditions. Using a prospectively collected CSF biobank, we analyzed 125 samples spanning multiple pathologies, including breast and lung LMD, glioblastoma, traumatic brain injury, and non-cancer neurological controls. Through optimized RNA extraction, library preparation, and deep sequencing, CSF-Seq generated robust and reproducible transcriptome-wide profiles despite the low abundance and fragmentation of cfRNA in CSF. CSF transcriptomes exhibited disease-specific expression, separating LMD from non-cancer controls and from non-LMD cancers, independent of CSF collection modality. Tumor-associated epithelial transcripts, including CEACAM6 and MUC1, were consistently enriched in LMD samples, whereas immune and CNS-associated transcripts were broadly detected across disease states, consistent with contributions from both tumor and non-tumor sources. Cross-site processing of matched samples demonstrated high concordance, indicating preservation of sample-specific transcriptional signatures across independent workflows. Importantly, we identified a collection method– independent LMD gene expression signature that was significantly associated with overall survival, supporting its potential prognostic relevance. Together, these findings establish CSF-Seq as a technically robust and clinically informative platform for transcriptomic biomarker discovery in CNS metastatic disease, offering a minimally invasive approach for disease characterization, risk stratification, and longitudinal monitoring in patients with LMD.

## INTRODUCTION

As advances in systemic cancer therapy extend patient survival, the incidence of brain metastases continues to rise, now affecting up to 30% of individuals with advanced cancer.^1^ These metastatic tumors are frequently therapy-resistant and are associated with profound neurological complications and poor survival.^2^ Among CNS metastatic diseases, leptomeningeal disease (LMD) represents the most aggressive and rapidly progressive manifestation, characterized by diffuse dissemination of tumor cells throughout the cerebrospinal fluid (CSF) and along the subarachnoid surfaces of the brain and spinal cord.^3^ LMD occurs most commonly in patients with breast and lung cancers, as well as melanoma and hematologic malignancies, and remains uniformly fatal with median survival measured in months.^4^ Current diagnostic approaches are limited: MRI lacks sensitivity for early or subtle disease, and CSF cytology— the clinical gold standard — fails to detect malignant cells in a substantial proportion of cases.^5,6^ As a result, diagnosis is often delayed, and clinical management is constrained by limited biological insight into disease burden and progression.

A major barrier to improving diagnosis and treatment is the limited access to tumor material from the CNS. Brain biopsy carries significant morbidity and is therefore rarely performed outside of diagnostic necessity, restricting opportunities for molecular characterization and biomarker discovery. Consequently, much of what is known about metastatic disease biology is inferred from primary tumors or extracranial metastases, despite growing evidence that tumor cells undergo extensive adaptation within the CNS microenvironment.^7,8^ This disconnect between the site of disease, and the source of available molecular data has constrained both biological understanding and translational progress.

CSF provides a unique and underutilized biospecimen for molecular interrogation of CNS disease. Circulating throughout the ventricular system and subarachnoid space, CSF directly interfaces with both normal brain tissue and metastatic lesions, carrying tumor-derived nucleic acids, immune signals, and cellular debris.^9^ Prior studies have demonstrated the clinical utility of CSF-derived cell-free DNA (cfDNA),^10,11^ which can reveal tumor-specific mutations even when not detected in plasma.^12^ However, while cfDNA is well suited for detecting genetic alterations, it offers limited insight into active transcriptional programs, tumor–microenvironment interactions, and dynamic disease states. In contrast, cell-free RNA (cfRNA) can capture gene expression from tumor cells, immune populations, and resident CNS cells, providing a functional readout of disease biology and therapeutic response. Despite this potential, transcriptomic profiling of CSF has remained technically challenging due to low RNA abundance, RNA fragmentation, and susceptibility to degradation. As a result, most prior studies have relied on targeted assays that assess a limited number of transcripts,^13^ restricting discovery of broader transcriptional programs and novel biomarkers.

To overcome these challenges, we developed CSF-Seq, a method for whole-transcriptome sequencing of cell-free RNA (cfRNA) from human CSF. Through optimized RNA extraction, library preparation, and deep sequencing, CSF-Seq enables reproducible profiling of both coding and noncoding transcripts from low-input samples. In this study, we applied CSF-Seq to a large cohort of CSF specimens collected from patient with diverse neurological and oncologic conditions. We show that CSF transcriptomes capture both tumor-associated and CNS-derived gene expression signals, distinguish disease states, and remain stable across variation in collection modality, short-term handling, and independent processing sites. Importantly, we identify an LMD-associated gene expression signature that is independent of sampling method and significantly associated with patient survival, suggesting potential utility for patient risk stratification when used alongside existing clinical assessments. Together, these findings establish CSF-Seq as a scalable and clinically relevant platform for molecular profiling of CNS metastatic disease, providing a minimally invasive approach to characterize disease biology, stratify patient risk, and enable longitudinal monitoring in patients with LMD.

## RESULTS

### CSF-Seq enables whole-transcriptome profiling and integrated clinical analysis of patient CSF

Figure 1 summarizes the study design, CSF acquisition, CSF-Seq workflow, and downstream analytical framework. CSF samples were obtained using three clinical collection modalities — direct ventricular access (including via Ommaya reservoir), lumbar puncture, and craniotomy-associated sampling (**Fig. 1a**)— reflecting the spectrum of clinical procedures used in routine care. Following collection, samples were processed to isolate cell-free nucleic acids, subjected to RNA extraction and cDNA synthesis, and prepared for whole-transcriptome sequencing (**Fig. 1b**). Libraries were sequenced to a target depth of approximately 1 x 10^8^ reads per sample, enabling broad transcriptome coverage despite the low abundance and fragmentation of CSF-derived RNA. This workflow establishes a standardized, high-depth sequencing pipeline compatible with clinically obtained CSF specimens.

**Figure 1.**
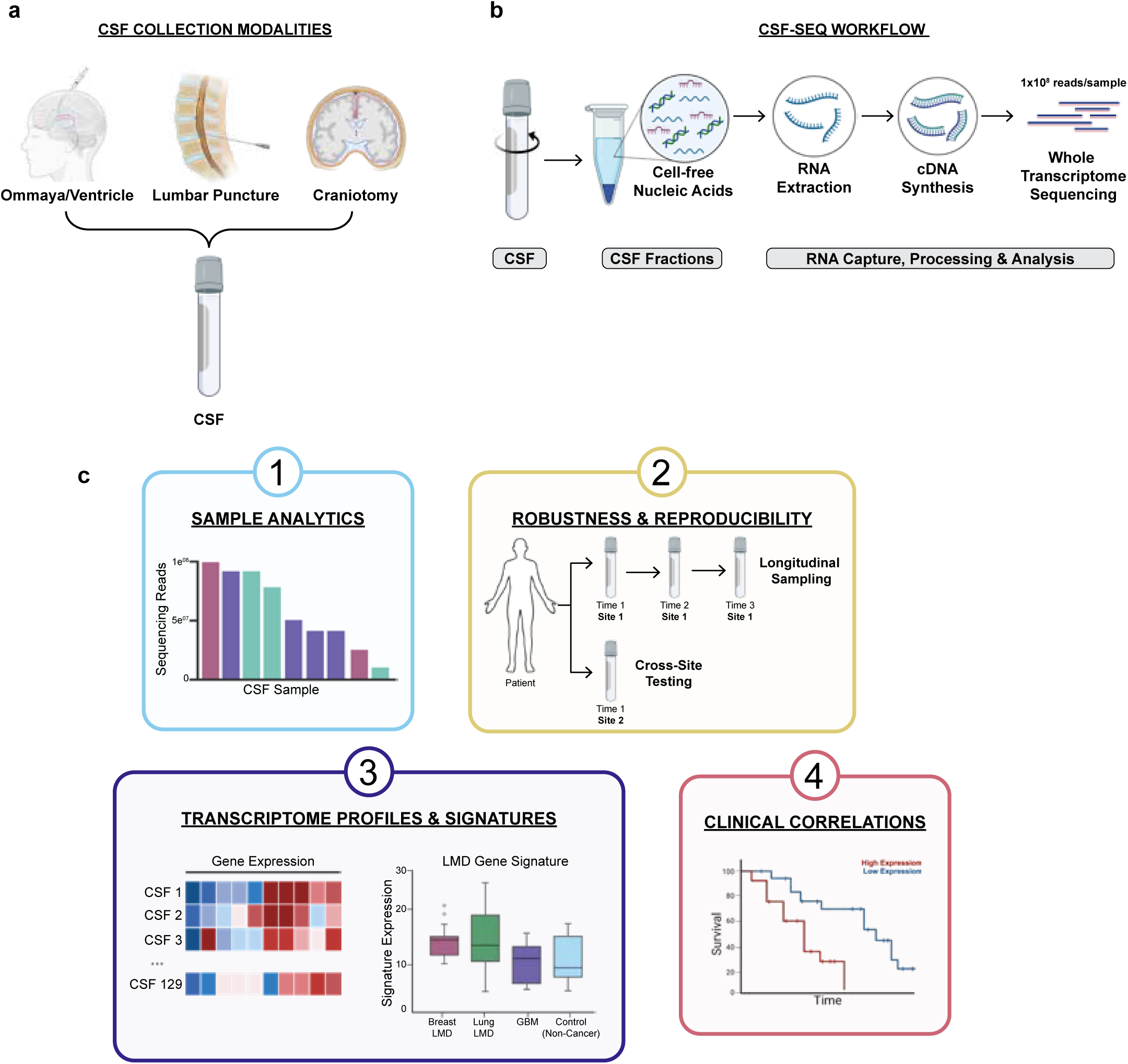
CSF-Seq enables whole-transcriptome profiling and integrated clinical analysis of patient CSF. **(a)** Schematic of CSF collection modalities used in this study, including ventricular access (via Ommaya reservoir), lumbar puncture, and craniotomy-associated sampling. **(b)** Overview of the CSF-Seq laboratory workflow. Following clinical acquisition, CSF undergoes processing for isolation of cell-free nucleic acids, RNA extraction, cDNA synthesis, and library preparation for whole-transcriptome sequencing. Libraries are sequenced to a target depth of ∼1×10^8^ reads per sample. **(c)** Schematic of the downstream analytical framework, organized into four modules: (1) sample-level sequence performance metrics, (2) assessments of technical robustness and reproducibility, (3) transcriptome-wide profiling and signature derivation, and (4) integration of molecular features with clinical outcome analyses. **CSF**, cerebrospinal fluid; **GBM**, glioblastoma; **LMD**, leptomeningeal disease.

Downstream analyses were organized into four complementary modules (**Fig. 1c**): (1) sample-level quality and sequencing performance metrics, (2) assessments of technical robustness and reproducibility across handling conditions, longitudinal sampling, and independent processing sites, (3) transcriptome-wide profiling and disease-associated signature discovery, and (4) integration of molecular features with clinical outcome data. Together, this framework enables evaluation of technical performance, biological signal, and clinical relevance within a single analytical pipeline. Applying this analytical framework to our clinical cohort, we show that CSF-Seq generates high-fidelity transcriptomic profiles that are reproducible across disease states, preserve biologically meaningful cell-type– associated signals, and remain robust to variability in clinical sampling conditions.

### CSF-Seq produces robust transcriptomic profiles across neurological and oncologic CSF samples

To evaluate the robustness of CSF-Seq data across disease contexts, we analyzed CSF samples spanning multiple clinical groups, including LMD from breast cancer (n = 16) and lung cancer (n = 22), glioblastoma (GBM; n = 15), traumatic brain injury (TBI; n = 42), non-cancer neurological controls (n = 30), and RNA from healthy brain tissue (n = 4) as a tissue control (**Fig. 2a**). Across all CSF samples, whole transcriptome sequencing produced high total read counts and substantial numbers of uniquely mapped reads (**Fig. 2b-c; Supplementary Fig. 1a**). While overall sequencing depth did not differ significantly between disease groups (**Fig. 2d**), modest but statistically significant differences were observed in uniquely mapped reads across disease groups (**Fig. 2e**).

**Figure 2.**
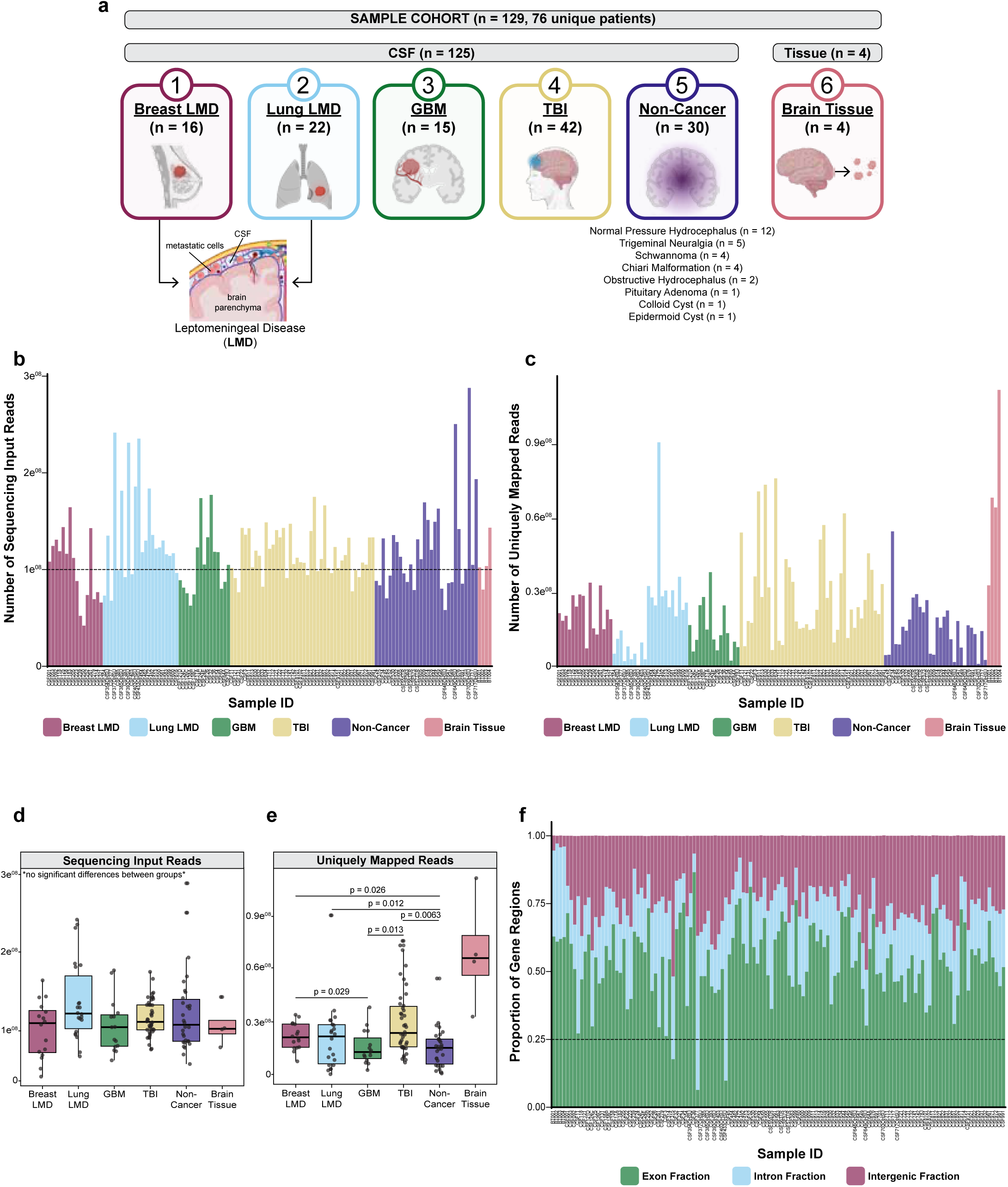
CSF-Seq generates consistent sequencing performance across neurological and oncologic CSF samples. **(a)** Overview of the study cohort. A total of n = 129 samples (76 unique patients) were analyzed, including CSF from breast LMD (n = 16), lung LMD (n = 22), glioblastoma (GBM; n = 15), traumatic brain injury (TBI; n = 42), non-cancer neurological conditions (n = 30), and RNA from brain tissue (n = 4). Total CSF samples: n = 125; tissue controls n = 4. **(b-c)** Sequencing performance across individual samples. Bar plots show **(b)** total input reads and **(c)** uniquely mapped reads per sample; each bar represents one CSF sample. **(d-e)** Group-level comparisons of sequencing metrics across disease categories. Boxplots show **(d)** total input reads and **(e)** uniquely mapped reads per sample; each dot represents one CSF sample. P values were calculated using two-sided Kruskal–Wallis tests with Dunn correction for multiple comparisons. **(f)** Distribution of reads across genomic features. Stacked bar plots show the proportion of exonic, intronic, and intergenic reads per sample; each bar represents one CSF sample. For boxplots, the center line denotes the median, box edges represent the 25^th^ and 75th percentiles, and whiskers extend to ±1.5× interquartile range (IQR). **CSF**, cerebrospinal fluid; **GBM**, glioblastoma; **LMD**, leptomeningeal disease; **TBI**, traumatic brain injury.

Analysis of read distribution across genomic features demonstrated that CSF-derived libraries contained consistent proportions of exonic, intronic, and intergenic reads across samples (**Fig. 2f**), supporting uniform library complexity and transcriptome representation. The relative fractions of gene regions were stable across disease categories (**Supplementary Fig. 1b-f**), indicating that CSF-Seq captures a reproducible spectrum of RNA species rather than being dominated by degraded or nonspecific background RNA.

Before downstream transcriptomic analyses, samples were subjected to multi-parameter quality filtering based on library size, exon fraction, and number of detected genes (**Supplementary Fig. 2a,b**). Independent QC metrics identified a small subset of samples with globally low transcript detection, which clustered together by low overall expression across protein-coding genes (**Supplementary Fig. 2c**). Overlap between orthogonal filtering criteria provided confidence in exclusion of truly low-quality libraries (**Supplementary Fig. 2d**). In total, 6 samples (4.7% of the cohort) were removed prior to normalization and biological analyses (**Supplementary Fig. 2e**), yielding a high-quality dataset for subsequent transcriptome-wide comparisons.

### CSF transcriptomes reflect disease-specific and cell-type–specific biological signals

Using the filtered high-quality dataset, we next assessed whether CSF-Seq profiles retain biologically informative gene expression patterns across disease states. The data processing and normalization workflow is summarized in Fig. 3a and included removal of non-expressed genes, restriction to protein-coding transcripts, and DESeq2-based normalization, yielding 18,649 genes for downstream analyses. Evaluation of gene counts before and after normalization (**Supplementary Fig. 3a**), and housekeeping gene behavior (**Supplementary Fig. 3b-g**) demonstrated stable expression across samples and disease groups, supporting the robustness of normalization procedures.

**Figure 3.**
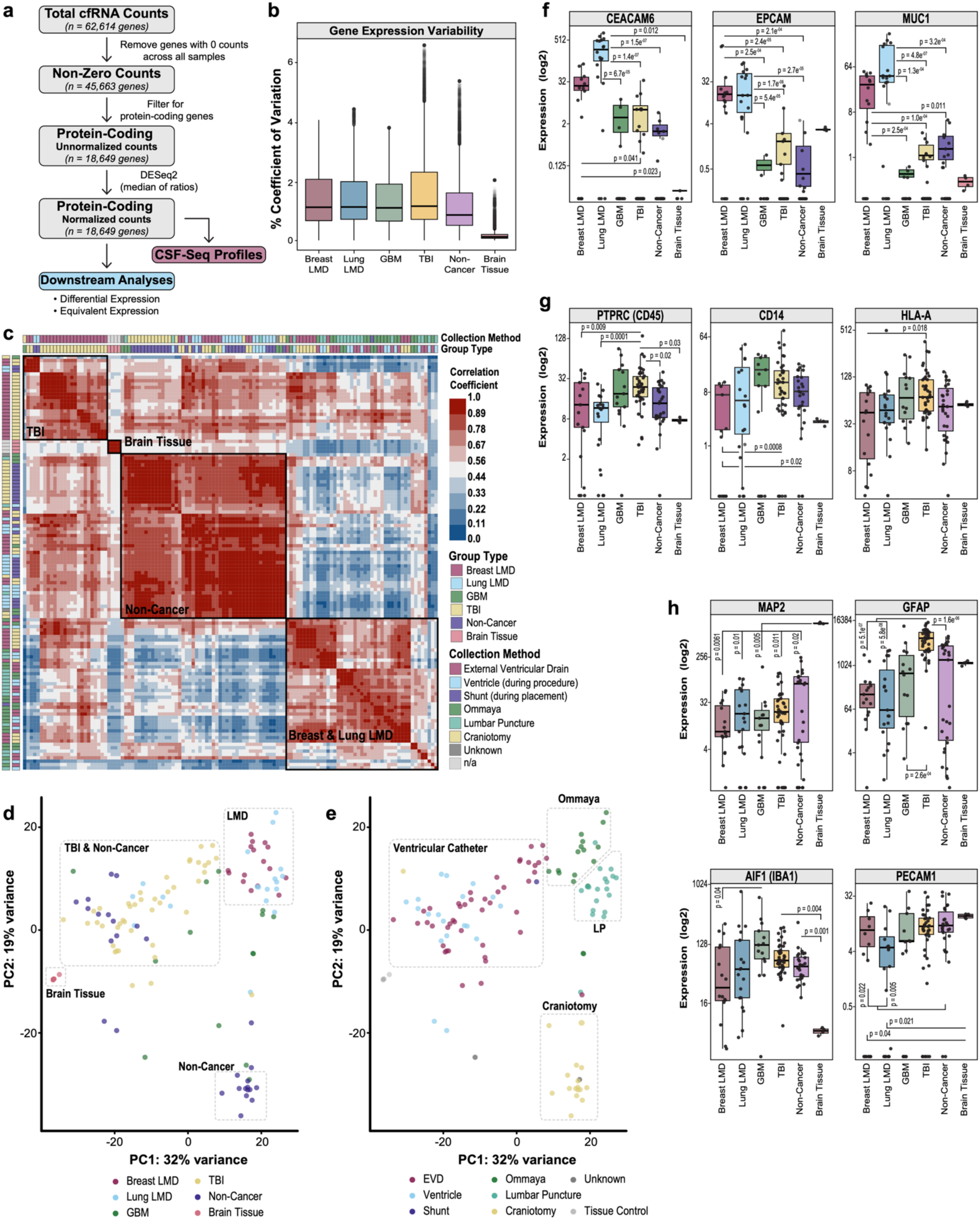
CSF transcriptomes reflect disease- and cell-type–specific biological signals. **(a)** Schematic of the CSF-Seq data processing workflow. Raw cfRNA counts (n = 62,614 genes) were filtered to remove genes with zero counts across all samples, restricted to protein-coding transcripts, and normalized using DESeq2 (median-of-ratios method), yielding 18,649 protein-coding genes for downstream analyses. **(b)** Gene expression variability across disease groups. Boxplot shows the distribution of coefficients of variation (CV) calculated for each protein-coding gene (n = 18,649 genes) using normalized expression values across samples within each disease category (breast LMD, n = 16; lung LMD, n = 19; GBM, n = 14; TBI, n = 42; non-cancer, n = 28; brain tissue, n = 4; total n = 123 samples); each dot represents one gene. **(c)** Pairwise sample correlation heatmap using normalized protein-coding gene expression (n = 18,649 genes; n = 123 samples). Color scale indicates Pearson correlation coefficient. Samples are annotated by disease group and collection modality. **(d-e)** Principal component analysis (PCA) of normalized protein-coding gene expression (n = 18,649 genes; n = 123 samples); each dot represents one CSF sample. **(d)** Samples annotated by disease group. **(e)** Samples annotated by collection modality. PC1 explains 32% of the variance and PC2 explains 19% of the variance. **(f-h)** Expression of representative cell-type–associated genes across disease groups. Boxplots show log2 normalized gene expression values; each dot represents one CSF sample (n = 123). **(f)** Tumor-associated genes (CEACAM6, EPCAM, MUC1). **(g)** Immune-associated genes (PTPRC/CD45, CD14, HLA-A). **(h)** Brain-resident cell markers (MAP2, GFAP, AIF1/IBA1, PECAM1). P values were calculated using two-sided Kruskal–Wallis tests with Dunn correction for multiple comparisons. For boxplots, the center line denotes the median, box edges represent the 25^th^ and 75th percentiles, and whiskers extend to ±1.5× interquartile range (IQR). **CV**, coefficient of variation; **EVD**, external ventricular drain; **GBM**, glioblastoma; **LMD**, leptomeningeal disease; **PCA**, principal component analysis; **TBI**, traumatic brain injury.

Gene expression variability across samples was evaluated using coefficients of variation for protein-coding genes (**Fig. 3b**). CSF samples from all disease groups demonstrated comparable variability, whereas brain tissue controls showed lower overall variability, consistent with higher RNA abundance and more homogeneous cellular composition in tissue relative to cell-free CSF. To evaluate similarity of transcriptomic profiles across samples, pairwise correlation analysis was performed using normalized gene expression values (**Fig. 3c**). This analysis revealed higher similarity among samples from the same disease category, with breast and lung LMD samples grouping separately from TBI and non-cancer CSF samples. Brain tissue samples formed an independent cluster distinct from all CSF samples, reflecting their fundamentally different composition compared to the cell-free CSF environment.

Principal component analysis further demonstrated separation of samples by clinical group (**Fig. 3d**). LMD samples segregated from TBI and non-cancer CSF along principal components explaining the largest proportion of variance, while brain tissue samples occupied a distinct region of transcriptional space. When samples were annotated by collection modality, partial clustering by sampling route was observed, with ventricular, Ommaya/lumbar puncture, and craniotomy-derived samples forming separable groups (**Fig. 3e**). However, disease-associated separation remained evident across collection methods, indicating that transcriptomic differences were not solely driven by sampling route. Additional PCA analyses stratified by clinical subgroups, technical variables, and matched sampling further confirmed that clustering patterns were driven by disease state rather than batch effects or personnel (**Supplementary Fig. 4**). Similar patterns were observed for gene expression variability (**Supplementary Fig. 5a,b**), housekeeping gene behavior (**Supplementary Fig. 5c,d**), and unsupervised clustering (**Supplementary Fig. 5e,f**), when analyses were restricted to cross-sectional samples (one sample per patient), indicating that variability metrics and global transcriptome profiles were not driven by repeated sampling or longitudinal collections.

To determine whether disease-associated clustering reflected biologically relevant cell-type signals, we examined expression of representative epithelial, immune, and brain-resident cell markers across sample groups (**Fig. 3f–h**). Epithelial and tumor-associated genes, including CEACAM6, EPCAM, and MUC1, were significantly enriched in CSF from breast and lung LMD samples relative to GBM, TBI, and non-cancer CSF, as well as relative to brain tissue controls (**Fig. 3f**). In contrast, immune-associated genes such as PTPRC (CD45), CD14, and HLA-A were broadly expressed across all CSF disease groups, consistent with the presence of immune-derived RNA in CSF regardless of etiology (**Fig. 3g**). Markers of brain-resident cell populations were readily detected in CSF and exhibited distinct patterns across disease states (**Fig. 3h**). The neuronal marker MAP2 was highest in brain tissue controls, whereas astrocytic (GFAP) and microglial (AIF1/IBA1) markers were broadly detected across CSF samples, with higher expression in conditions associated with neuroinflammation and tissue injury. Endothelial-associated gene expression (PECAM1) was detected across both CSF and tissue samples, indicating that vascular-associated transcripts contribute to the CSF cell-free RNA pool. Together, these results suggest that CSF-Seq captures signals from multiple cellular compartments, including tumor-associated, immune, and CNS-resident sources. To ensure that observed disease-associated transcriptional patterns were not driven by pre-analytical or technical variability, we next evaluated the robustness and reproducibility of CSF-Seq across handling conditions, longitudinal sampling, and inter-site processing.

### CSF-Seq is robust to variation in sample handling and processing across sites

To evaluate the technical robustness of the CSF-Seq workflow, we assessed the impact of short-term handling conditions and longitudinal sampling on transcriptomic profiles (**Fig. 4a–c**). Pre-analytical variables included temperature during short-term processing (on ice versus room temperature) and duration of processing at room temperature (30 versus 60 minutes) prior to RNA extraction (**Fig. 4a,b**).

**Figure 4.**
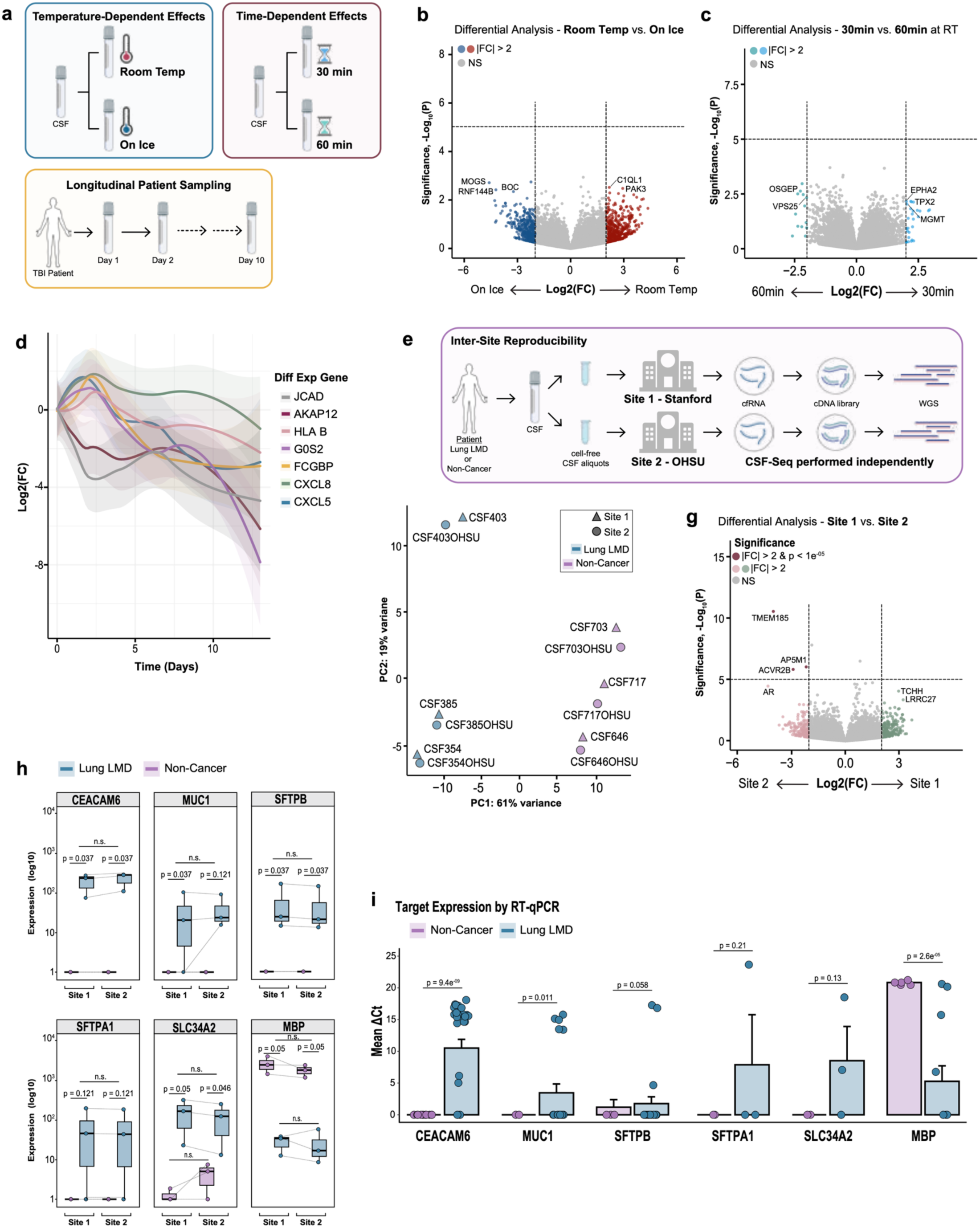
CSF-Seq is robust to variation in sample handling, longitudinal sampling, and inter-site processing. **(a)** Temperature-dependent handling effects. Schematic illustrating short-term CSF processing conditions (on ice, n = 4, versus room temperature, n = 4) followed by differential expression analysis using DESeq2. Volcano plot shows log2 fold change versus −log10(adjusted p value); each dot represents one protein-coding gene. Genes exceeding |log2 fold change| > 2 and adjusted p < 1 x 10^−5^ are colored; NS, not significant. **(b)** Processing time-dependent effects. Schematic illustrating comparison of 30 minutes (n = 11) versus 60 minutes (n = 11) at room temperature prior to RNA extraction. Volcano plot shows log2 fold change versus −log10(adjusted p value); each dot represents one protein-coding gene. Genes exceeding |log2FC| > 2 adjusted p < 1 x 10^−5^ are colored; NS, not significant. **(c)** Longitudinal sampling in TBI. Schematic illustrating serial CSF collection (days 1, 2, and 10). Line plot shows representative differentially expressed genes across time; curves represent LOESS-smoothed log2 fold change trajectories. Longitudinal CSF samples collected from the same patients at multiple time points were treated as repeated measures, with each time point representing a distinct biological sample. **(d)** Inter-site reproducibility workflow. Schematic illustrating independent processing of matched CSF aliquots at two institutions (Stanford and OHSU), including RNA extraction, cDNA synthesis, library preparation, and whole-transcriptome sequencing. **(e)** PCA of normalized protein-coding gene expression from matched inter-site samples (Site 1, n = 6; Site 2, n = 6; 3 lung LMD and 3 non-cancer samples per site); each dot represents one sample. PC1 explains 61% of the variance and PC2 explains 19% of the variance. Samples are annotated by processing site and disease group. **(f)** Differential expression analysis comparing processing Site 1 versus Site 2 using DESeq2. Volcano plot shows log2 fold change versus −log10(adjusted p value); each dot represents one protein-coding gene. Genes exceeding |log2FC| > 2 and adjusted p < 1×10⁻⁵ are colored; NS, not significant. **(g)** Expression of representative tumor-associated and brain-associated genes across matched inter-site samples. Boxplots show log10 normalized expression values for CEACAM6, MUC1, SFTPB, SFTPA1, SLC34A2, and MBP, stratified by site and disease group; each dot represents one CSF sample (Site 1, n = 6; Site 2, n = 6). P values were calculated using two-sided Wilcoxon signed-rank tests. **(h)** Target expression measured by RT–qPCR in lung LMD versus non-cancer control CSF. Bar plots show mean ΔCt values for indicated genes; error bars represent standard error of the mean (SEM). Each dot represents one biological CSF sample averaged across technical replicates. The number of biological samples analyzed varied by target gene (lung LMD, n = 3–22; non-cancer, n = 2–7). P values were calculated using two-sided Wilcoxon rank-sum test (Mann– Whitney U). For boxplots in (g), the center line denotes the median, box edges represent the 25th and 75th percentiles, and whiskers extend to ±1.5× interquartile range (IQR). **LMD**, leptomeningeal disease; **LOESS**, locally estimated scatterplot smoothing; **NS**, not significant; **RT–qPCR**, reverse transcription quantitative PCR; **TBI**, traumatic brain injury.

Differential expression analysis comparing samples maintained on ice versus room temperature did not identify transcripts meeting combined thresholds for statistical significance and large effect size, with most genes exhibiting modest fold changes and failing to reach significance (**Fig. 4a**). Likewise, comparison of samples processed after 30 minutes versus 60 minutes at room temperature revealed no statistically significant differences in gene expression, and transcript abundance was highly concordant across conditions (**Fig. 4b**). These results indicate that moderate variation in short-term handling temperature and processing time does not substantially perturb CSF transcriptomic profiles captured by CSF-Seq.

To assess the ability of CSF-Seq to capture temporal transcriptional dynamics, we analyzed longitudinal samples collected from patients with traumatic brain injury (TBI) across multiple time points (days 1, 2, and 10) (**Fig. 4c**). This analysis identified subsets of transcripts exhibiting marked changes over time, consistent with evolving injury-associated and inflammatory responses, while many genes showed stable expression across the same interval (**Supplementary Fig. 6a**). Although longitudinal sampling in breast LMD and GBM was limited (n = 2 patients per group), heatmap-based and gene-level analyses revealed transcripts exhibiting consistent directional changes across patients between time points (**Supplementary Fig. 6b–e**), supporting the ability of CSF-Seq to capture temporal transcriptional dynamics in multiple disease settings.

To evaluate inter-site reproducibility, matched aliquots of the same CSF samples from patients with lung LMD or non-cancer neurological conditions, specifically normal pressure hydrocephalus, were processed independently at two institutions using separate workflows spanning RNA extraction, library preparation, and sequencing, with the original CSF specimen serving as the only shared input (**Fig. 4d**). Principal component analysis demonstrated that matched samples processed at different sites clustered closely together, with sample identity outweighing site of processing as a determinant of transcriptional similarity (**Fig. 4e**), indicating strong preservation of sample-specific transcriptional signatures across independent workflows. Direct comparison of gene expression between sites revealed high concordance, with only three protein-coding genes meeting criteria for differential expression (**Fig. 4f**). This minimal number of site-associated differences suggests that independent processing workflows introduce little systematic bias into CSF-Seq profiles.

To further assess reproducibility of biologically relevant tumor-associated transcripts, expression of established lung cancer and epithelial markers, including CEACAM6, MUC1, SFTPB, SFTPA1, and SLC34A2, as well as the brain-associated gene MBP, was examined across matched site pairs (**Fig. 4g; Supplementary Fig. 6f,g**). These genes displayed consistent relative expression patterns between lung LMD and non-cancer samples at both sites, with no significant differences attributable to processing location. Concordant results were obtained using orthogonal RT– qPCR assays (**Fig. 4h; Supplementary Fig. 6h**), further supporting the quantitative reliability of CSF-Seq–based detection of disease-associated transcripts.

Collectively, these analyses demonstrate that CSF-Seq provides stable and reproducible transcriptomic profiles across variations in short-term sample handling, longitudinal sampling, and independent laboratory processing, while preserving biologically meaningful disease-associated transcriptional signals.

### CSF-Seq identifies a collection-independent LMD gene signature associated with patient survival

To identify transcriptional programs associated with leptomeningeal disease (LMD), we compared CSF samples from patients with LMD to non-cancer neurological controls while accounting for CSF collection modality. Differential expression analysis revealed consistent upregulation of epithelial and tumor-associated transcripts in both breast and lung LMD samples relative to non-cancer CSF across collection methods (**Fig. 5a,b**), including CEACAM6, MUC1, EPCAM, KRT7, and KRT19. When stratified by primary tumor type, breast- and lung-specific LMD comparisons revealed a 30% shared transcriptional core (1,342 overlapping differentially expressed genes), alongside partially distinct gene sets specific to each tumor type, indicating the presence of both conserved and tumor type-specific transcriptional programs in LMD CSF (**Supplementary Fig. 7a**).

**Figure 5.**
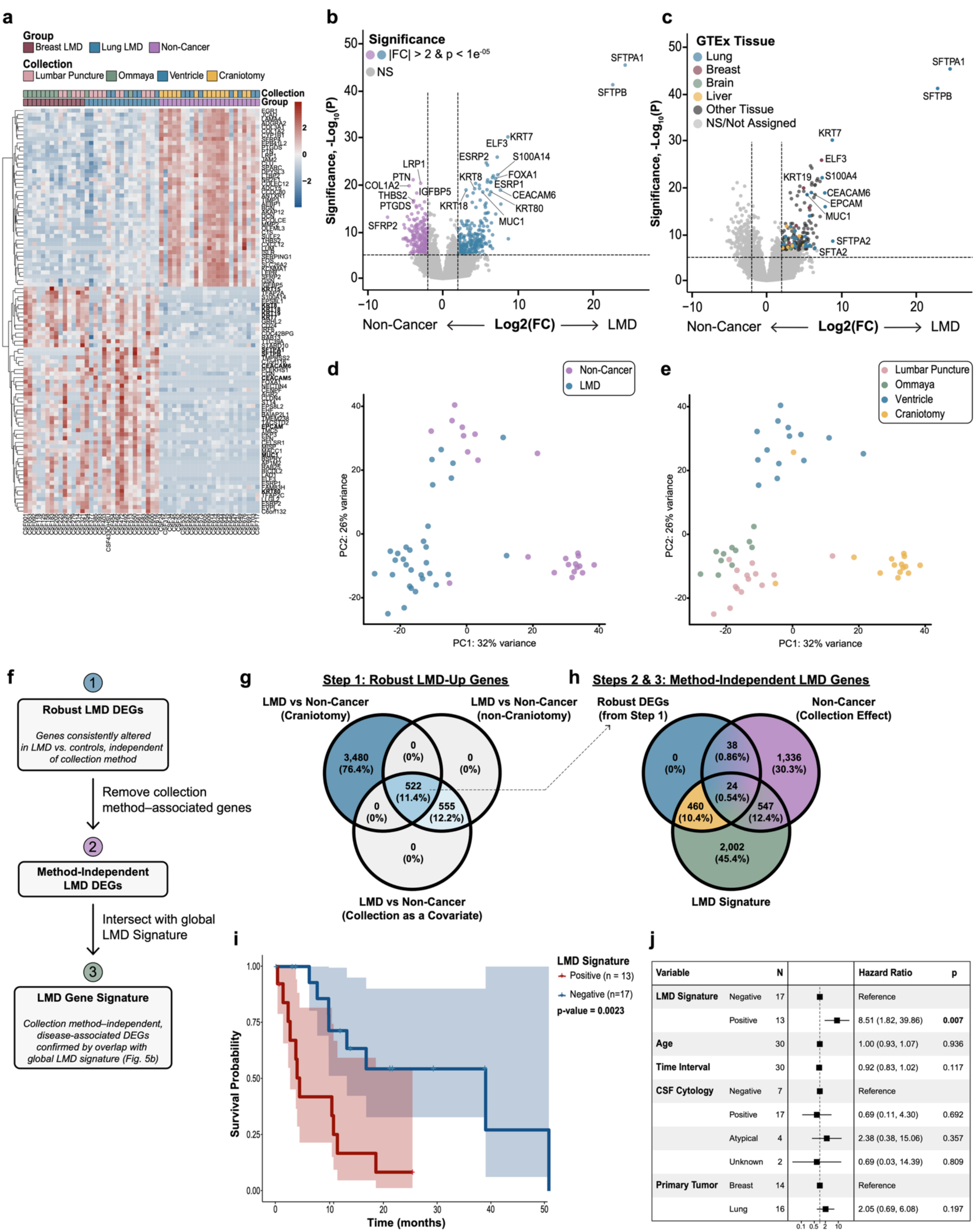
Derivation of a collection-independent LMD transcriptional signature associated with patient survival. **(a)** Heatmap of differentially expressed genes across LMD and non-cancer CSF samples (n = 54). Gene expression values are scaled by row. Samples are hierarchically clustered and annotated by disease group and collection modality. **(b)** Differential expression analysis comparing LMD versus non-cancer controls. Volcano plot shows log2 fold change versus −log10(adjusted p value); each dot represents one protein-coding gene. Genes exceeding |log2FC| > 2 and adjusted p < 1×10⁻⁵ are colored. **(c)** Volcano plot as in **(b)**, annotated by GTEx tissue enrichment category for significantly upregulated genes. (**d-e**) PCA of normalized protein-coding gene expression from LMD and non-cancer samples (n = 54); each dot represents one CSF sample. **(d)** Samples annotated by disease group. **(e)** Samples annotated by collection modality. **(f)** Schematic of multistep filtering workflow used to derive the collection-independent LMD signature. **(g)** Venn diagram showing overlap of significantly differentially expressed genes identified across three differential expression analyses (using DESeq2) stratified by collection modality. **(h)** Venn diagram showing intersection of genes retained after modality-associated gene removal and global LMD differential expression analysis, yielding the final 460-gene collection-independent signature. **(i)** Kaplan–Meier survival plot stratifying LMD patients by high versus low LMD signature expression (n = 30 patients). P value calculated using two-sided log-rank test. **(j)** Forest plot of multivariable Cox proportional hazards model including LMD signature score, age, primary tumor type, CSF cytology status, and time from diagnosis to CSF collection. Hazard ratios (HR) with 95% confidence intervals (CI) are shown. **GTEx**, Genotype-Tissue Expression; **LMD**, leptomeningeal disease; **PCA**, principal component analysis.

To further characterize the tissue-associated expression programs represented among LMD-associated transcripts, differentially expressed genes were annotated using Human Adult Genotype Tissue Expression (GTEx)^14^ reference profiles. Genes upregulated in LMD were preferentially associated with epithelial tissue expression signatures, including breast, lung, and liver, whereas comparatively fewer transcripts mapped to brain-enriched expression programs (**Fig. 5c**). Projection of breast and lung LMD gene sets onto GTEx tissue and tissue composition reference datasets demonstrated enrichment of epithelial-dominated tissue profiles rather than endothelial, fibroblast, adipose, or muscle-associated programs (**Supplementary Fig. 7b,c**). Consistent with this observation, projection of breast- and lung-derived LMD expression profiles into an independent brain metastasis dataset^15^ revealed selective enrichment in metastatic tumor cells (**Supplementary Fig. 7d,e**). Deconvolution of breast LMD CSF samples using a single-cell breast LMD reference^16^ further indicated enrichment of epithelial- and innate/adaptive immune-associated programs, consistent with known features of LMD biology (**Supplementary Fig. 7f-h**).

However, despite these consistent disease-associated signals, CSF collection modality substantially influenced gene expression patterns in LMD versus non-cancer controls (**Fig. 5d,e; Supplementary Fig. 8a–d**), motivating the development of a collection-independent transcriptional signature for LMD that would remain robust across heterogeneous clinical sampling conditions. Rather than restricting analyses to a single control subgroup, which would limit generalizability and reduce available sample size, we implemented a multistep filtering strategy to identify disease-associated transcripts consistently altered in LMD irrespective of control composition or collection route (**Fig. 5f**). At each step, both up- and down-regulated genes were retained, enabling an unbiased representation of LMD-associated transcriptional changes. First, we performed three independent differential expression analyses using DESeq2: (i) LMD versus non-cancer controls collected via craniotomy, (ii) LMD versus non-cancer controls collected via non-craniotomy routes (i.e., lumbar puncture, Ommaya, ventricular catheter), and (iii) LMD versus all non-cancer controls with collection modality included as a covariate in the DESeq2 model (**Supplementary Fig. 8e–g**). Genes significantly differentially expressed in LMD relative to controls were identified in each comparison, and only those consistently altered in the same direction across all three analyses were retained. This intersection yielded 522 LMD-associated genes (**Fig. 5g**), ensuring robustness to control composition and sampling route. Second, to exclude transcripts driven by sampling-related anatomical or technical effects independent of disease status, we performed differential expression analysis between craniotomy and non-craniotomy non-cancer controls (**Supplementary Fig. 8h**). Genes significantly associated with collection modality in the absence of malignancy were removed from the 522 LMD-associated genes previously identified, as these transcripts — particularly those enriched in brain tissue — were more sensitive to sampling location rather than disease biology. Third, the remaining genes were intersected with the global LMD versus non-cancer differential expression signature derived from the full cohort from Fig. 5b, ensuring concordance with disease-associated expression patterns observed at the cohort level rather than subset-specific contrasts. This final filtering step yielded a collection method–independent LMD transcriptional signature consisting of 460 genes (**Fig. 5h**).

To further illustrate the robustness of this signature, expression of representative genes consistently increased or decreased in LMD across all filtering steps was visualized across LMD samples and non-cancer controls stratified by collection modality (**Supplementary Fig. 8i**), demonstrating preserved disease-associated directionality independent of control sampling route. Gene set enrichment analysis performed separately on genes up- and down-regulated in LMD revealed distinct biological programs (**Supplementary Fig. 8j,k**). Genes consistently up-regulated in LMD were enriched for cell cycle, chromosome segregation, spindle assembly, and epithelial differentiation pathways, consistent with a highly proliferative tumor-associated transcriptional state. In contrast, genes consistently down-regulated in LMD were enriched for extracellular matrix organization, collagen-containing matrix, growth factor binding, and glial projection, suggesting loss of normal stromal and glial-associated programs in CSF from LMD patients.

To assess clinical relevance, signature scores were calculated for individual patients and used to stratify LMD cases into high- and low-signature expression groups. Kaplan–Meier analysis demonstrated significantly reduced overall survival among patients with high LMD signature expression (**Fig. 5i**; p = 0.0023). In multivariable Cox proportional hazards modeling, high LMD signature expression remained significantly associated with increased hazard of death (hazard ratio = 8.51, 95% CI = 1.82–39.86, p = 0.007) after adjustment for clinical covariates, whereas age, primary tumor type, CSF cytology status, and time from diagnosis to CSF collection were not associated with outcome (**Fig. 5j**). These results indicate that CSF-derived transcriptional signatures provide prognostic information beyond standard clinical parameters in patients with LMD.

Finally, to evaluate whether CSF-Seq resolves distinct disease-specific transcriptional programs across diverse neurological pathologies beyond LMD, differential expression and gene set enrichment analyses were performed across glioblastoma (GBM) (**Supplementary Fig. 9a,b**) and traumatic brain injury (TBI) (**Supplementary Fig. 9c,d**) CSF samples. Each disease groups exhibited largely non-overlapping differentially expressed gene sets, confirming that CSF-Seq distinguishes distinct pathological states (**Supplementary Fig. 9e,f**). In contrast, genes not differentially expressed across disease groups were enriched for pathways related to mitochondrial respiration, proteostasis, autophagy, and vesicle trafficking, suggesting the presence of a conserved CSF background program reflecting universal mechanisms of cellular maintenance and stress adaptation that remain independent of disease-specific transcriptional states (**Supplemental Fig. 9g,h**).

Together, these analyses demonstrate that CSF-Seq identifies biologically coherent, collection-independent transcriptional signatures in LMD that are reproducible across sampling routes, supported by concordant enrichment in external reference datasets, and strongly associated with patient outcomes.

## DISCUSSION

Leptomeningeal disease (LMD) remains one of the most devastating complications of systemic cancer, with limited tools for early detection, biological characterization, and prognostication. Current clinical assessment relies primarily on neuroimaging and CSF cytology, both of which have limited sensitivity and provide little insight into underlying disease biology.^3,17^ In this study, we demonstrate that CSF-Seq enables robust, reproducible transcriptomic profiling of CSF-derived cell-free RNA across heterogeneous clinical sampling conditions and disease states, and that these profiles capture biologically and clinically meaningful programs associated with patient outcomes. Together, these findings establish CSF transcriptomics as a feasible and informative approach for molecular interrogation of LMD and other CNS pathologies.

A central contribution of this work is the identification of a collection method–independent LMD gene signature that is strongly associated with overall survival. Rather than reflecting a single biological axis, the 460-gene signature appears to represent a composite transcriptional state, integrating information about tumor-associated proliferation, epithelial identity, and disruption of normal CNS-associated programs that may be missed by conventional diagnostic measures alone. Although CSF cytology and neuroimaging remain central to the clinical evaluation of LMD, both modalities are limited in sensitivity and provide minimal insight into underlying molecular programs.^3,5,18–21^ Notably, cytology status was not associated with survival in this cohort, a finding which has also been reported across other cohorts.^19^ By contrast, patients with high signature expression experienced significantly shorter time-to-death, and this association remained robust after adjustment for clinical covariates, including age, primary tumor type, CSF cytology status, and time from diagnosis to CSF collection, underscoring that CSF-derived transcriptomic profiling offers an orthogonal and biologically-rich measure of tumor state and microenvironmental disruption. While integrated MRI or ctDNA analyses were beyond the scope of this study, future work combining CSF transcriptomics with imaging and genomic and/or proteomic assays will be critical to defining how these modalities jointly inform disease burden, progression, and therapeutic response.

The multistep filtering strategy used to derive this signature addresses a key challenge in CSF-based biomarker discovery: the substantial influence of sampling route and control composition on gene expression profiles.^22–24^ Many prior CSF biomarker studies rely on narrowly matched control groups or single comparison frameworks, which can limit generalizability and inflate cohort-specific effects.^25–27^ Here, by explicitly modeling and filtering across multiple control groups and sampling routes, and by retaining both up- and down-regulated genes at each step, we prioritized transcriptional changes that were consistently associated with LMD irrespective of reference group or collection modality. This approach reduces susceptibility to confounding driven by technical factors or anatomical differences associated with CSF collection route and provides a conceptual framework for biomarker discovery in clinically heterogeneous biofluids.

Biologically, the LMD signature reveals a coordinated remodeling of the CSF transcriptome characterized by gain of tumor-associated epithelial and proliferative programs alongside loss of normal CNS-supportive signals. Genes consistently up-regulated in LMD were enriched for cell cycle progression, chromosome segregation, spindle assembly, and epithelial differentiation pathways, consistent with active tumor proliferation and structural adaptation within the leptomeningeal compartment.^28–30^ In contrast, genes consistently down-regulated in LMD were enriched for extracellular matrix organization, collagen-containing matrix, growth factor binding, and glial projection–associated terms. These patterns suggest erosion of stromal, glial, and extracellular support programs that are otherwise detectable in non-cancer CSF, reflecting replacement of normal CNS-associated transcriptional signals by tumor-dominated programs. Similar disruption of glial and extracellular matrix interactions has been described in brain metastases and other CNS malignancies.^31,32^

Together, these results point to several clinically relevant applications of CSF-Seq across the course of CNS metastatic disease. At diagnosis, transcriptomic profiling could complement imaging and cytology. Particularly in cases with negative or equivocal cytologic findings, detection of tumor-associated transcriptional programs may precede overt malignant cell detection.^4,7,11,16,21^ Risk stratification based on the LMD signature could help identify patients at highest risk of rapid progression, informing prognosis and clinical decision-making. Longitudinal CSF sampling further raises the possibility of monitoring disease dynamics or treatment response through changes in transcriptional state over time. Although longitudinal LMD sampling was limited in this study, proof-of-principle analyses and ongoing clinical trial efforts suggest that such applications may be feasible.^33,34^

Limitations should be considered when interpreting these findings and framing the significance of this work. First, while this study includes multiple disease groups, sampling routes, and independent processing sites, cohort sizes— particularly for longitudinal and subtype-specific analyses — remain modest. Larger, prospectively collected, multi-center cohorts will be required to refine prognostic models and evaluate assay performance across broader clinical contexts. Second, CSF-Seq profiles bulk cell-free RNA and therefore cannot directly resolve the cellular origins of individual transcripts. While prior work suggests that cellular CSF sequencing in LMD primarily captures tumor and immune cell populations,^16,35,36^ bulk sequencing of cell-free CSF RNA enables detection of transcriptional programs arising not only from circulating tumor and immune cells but also from the CNS microenvironment, including stromal and glial populations that are not physically present in the CSF cellular fraction. Integration with complementary approaches such as single-cell RNA sequencing, proteomics, or ctDNA profiling may help better resolve the cellular sources and biological context of these transcriptional programs. Finally, the association between the LMD signature and survival is observational. While the signature captures biologically coherent programs linked to aggressive disease, causality cannot be inferred from these data alone. Future functional and interventional studies will be required to determine whether these transcriptional states actively drive disease progression or reflect downstream consequences of advanced pathology.

Despite these limitations, this study advances the field in several important ways. We demonstrate that CSF-Seq is technically robust, reproducible across clinical and laboratory variability, and capable of resolving disease-specific transcriptional programs across neurological and oncologic conditions. We introduce a principled, multistep strategy for deriving collection-independent molecular signatures in heterogeneous clinical biofluids. Most importantly, we identify a CSF-Seq-derived LMD signature that captures clinically meaningful disease biology and stratifies patient outcomes beyond standard clinical metrics. Together, these findings position CSF transcriptomics as a promising minimally invasive tool for advancing diagnosis, prognostication, and biological understanding of LMD and other CNS pathologies.

## METHODS

### Human biospecimen collection

Cerebrospinal fluid (CSF) samples were obtained from patients receiving clinical care at Stanford Hospital under protocols approved by the Stanford Institutional Review Board. Additional GBM CSF samples were obtained from Mayo Clinic under Institutional Review Board-approved protocols (NCT04692337, NCT04692324, or the Mayo Clinic neuro-oncology biorepository). The study complied with HIPAA regulations. Written informed consent permitting research use of biospecimens and access to clinical data was obtained from all participants. Samples were not de-identified prior to analysis; specimens shared or obtained externally were de-identified. A total of 125 CSF samples were collected from 75 patients, including individuals with leptomeningeal disease (LMD) from primary lung or breast cancer, glioblastoma (GBM), traumatic brain injury (TBI), and non-cancer neurological controls. CSF was collected via direct ventricular access (including Ommaya reservoir, external ventricular drain, or ventricular shunt), lumbar puncture, or cisternal drainage during intraoperative craniotomy. Only CSF volumes exceeding those required for clinical diagnostics were used for molecular profiling.

### CSF processing and storage

Following collection, CSF samples were immediately placed on ice and centrifuged to separate cellular and cell-free fractions (1,000 x g for 10 minutes at 4°C for Stanford samples; 400 x g for 10 minutes at 4°C for Mayo GBM samples). Supernatant was aliquoted and stored at –80°C until RNA extraction. Clinical metadata including diagnosis, collection modality, and sample volume were recorded in a centralized biorepository database.

### Cell-free RNA extraction

Cell-free RNA (cfRNA) was isolated from 1–2 mL CSF using the Plasma/Serum Circulating and Exosomal RNA Purification Mini Kit (Norgen Biotek, #51000, #29500). Residual DNA contamination was removed using Baseline-ZERO DNase (Lucigen, #DB0715K). RNA was purified and concentrated using RNA Clean & Concentrator-25 (Zymo Research, #R1017, #R1018) and eluted in 18 µL nuclease-free water. RNA concentration and purity were assessed by spectrophotometry (NanoDrop, Thermo Scientific) using 2 µL of elution.

RNA was stored at –80 °C in aliquots (2 x 5 µL, 3 x 2 µL).

### Library preparation and sequencing

Strand-specific sequencing libraries were prepared using the SMARTer Stranded Total RNA-Seq Kit v3 - Pico Input Mammalian (Takara Bio, #634486), which includes ribosomal RNA depletion. Libraries were purified using AMPure XP beads (Beckman Coulter, #A63880) and evaluated for size distribution and concentration using Agilent Bioanalyzer 2100 and spectrophotometry (NanoDrop). Libraries were sequenced on an Illumina NovaSeq 6000 platform using 2 × 150 bp paired-end reads to a target depth of approximately 1 × 10⁸ reads per sample.

### RNA-seq preprocessing pipeline

Raw sequencing reads (FASTQ) underwent adapter trimming and quality filtering using Sickle (v1.33) (paired-end mode).^37^ Read quality metrics were assessed using FastQC (v0.11.9).^38^ Contamination screening was performed using FastQ Screen (v0.14.1)^39^ against a multispecies reference genome database. Filtered reads were aligned to the human reference genome (GRCh38; GENCODE v34 annotation)^40^ using STAR (v2.7.10a)^41^ in two-pass mode. Duplicate reads were marked and alignment metrics (e.g., insert size, duplication rate) collected using Picard tools (v2.27.5). Aligned reads were coordinate-sorted using SAMtools (v1.9).^42^ Gene-level counts were generated using HTSeq (v0.13.5). All tools were run with default parameters unless otherwise specified. The computational pipeline was executed within a containerized environment.

### Gene and sample filtering

Gene annotations were obtained using biomaRt (v2.62). Analyses were restricted to protein-coding genes. Genes with zero counts across all samples were removed. After filtering and normalization, 18,649 genes were retained for downstream analyses. Quality control metric included library size, exon mapping fraction, and number of detected genes. Samples failing orthogonal quality control thresholds were excluded prior to normalization (n = 6 samples, 4.7%). No batch correction was applied. Principal component analysis was performed with samples annotated by sequencing run and processing personnel to assess potential batch effects.

### Differential Gene Expression Analysis

Differential expression analyses were performed using DESeq2 (v1.46). Raw counts were normalized using the median-of-ratios method. DESeq2-independent filtering was applied to remove low-count genes. Negative binomial generalized linear models were fit for each contrast, and gene-wide dispersions estimates were obtained using empirical Bayes shrinkage. Statistical significance was determined using two-sided Wald tests with Benjamini–Hochberg false discovery rate (FDR) correction. Genes with FDR-adjusted p < 0.05 and |log₂ fold change| > 2 were considered statistically significant unless otherwise specified. Standard differential expression models included disease group as the primary variable. Age and sex were not included in differential expression models. Collection modality and processing site were included as covariates only in analyses explicitly evaluating sampling or inter-site effects. For analyses involving repeated sampling, patient identity was included as a fixed effect in the DESeq2 design formula. For volcano plot visualizations, a more stringent threshold (adjusted p < 1 × 10⁻⁵ and |log₂ fold change| > 2) was used to highlight strongly differentially expressed genes for graphical emphasis. Longitudinal expression trajectories were visualized using LOESS regression with 95% confidence intervals.

### External dataset mapping

Reference tissue expression profiles were obtained from the Genotype-Tissue Expression (GTEx) database. GTEx expression matrices were transformed into gene-by-tissue format. For genes present in both GTEx and differential expression results, tissue enrichment was assigned based on maximum average expression across tissues. A predefined subset of tissues (lung, breast, liver, skeletal muscle, whole blood, and selected brain regions) was used for restricted tissue assignment. Genes were assigned to the tissue with highest expression exceeding a threshold expression value of 50 normalized units. Genes not exceeding this threshold were categorized as non-specific. Projection into external single-cell datasets was performed using publicly available leptomeningeal metastasis and brain metastasis reference datasets. Cellular deconvolution was performed using CIBERSORTx^43^ with B-mode batch correction. Reference single-cell RNA-seq data were obtained from Prakadan et al. (Nat Commun 2021; accession SCP1332, dbGaP phs002416.v1.p1),^16^ comprising adaptive immune, innate immune, and malignant cell populations. Signature matrices were generated from the reference dataset using default parameters.

### Gene Ontology enrichment analysis

Gene Ontology (GO) enrichment analysis was performed using the clusterProfiler R package (v4.14.6). Enrichment testing was conducted using over-representation analysis (ORA) implemented through the enrichGO function. Differentially expressed gene sets were compared against a background universe consisting of all tested protein-coding genes detected in the dataset. Enrichment was evaluated across the Biological Process, Molecular Function, and Cellular Component GO ontologies. For differential expression analyses performed across disease comparisons, enrichment analyses were conducted using combined sets of significantly differentially expressed genes regardless of directionality. For analyses involving the collection-independent leptomeningeal disease (LMD) transcriptional signature, enrichment analyses were performed separately for genes upregulated and downregulated in LMD. Statistical significance for enrichment was defined as a Benjamini–Hochberg false discovery rate (FDR) < 0.05.

### Survival and prognostic modeling

An LMD-associated gene expression signature was constructed using significantly differentially expressed genes from the LMD versus control comparison. Per-sample signature scores were calculated using the AddModuleScore function in Seurat (v5.2.1).^44^ Patients were stratified using a score threshold of zero. Overall survival was evaluated using Kaplan–Meier estimation with two-sided log-rank testing. Survival analyses were performed on patients with LMD for whom survival data were available (n = 30). Cox proportional hazards regression was performed using two-sided tests, adjusting for age, primary tumor type, cytology status, and time from diagnosis to CSF sampling. Model assumptions, including proportional hazards, linearity of continuous variables, and influence of outliers, were validated using Schoenfeld residuals.

### Reverse-transcription quantitative PCR (RT-qPCR)

Cell-free RNA was extracted and purified as described above. RT–qPCR was performed using the TaqMan RNA-to-Ct 1-Step Kit (Thermo Fisher Scientific, #4392938). Each 20-µL reaction contained 10 µL 2× RT–PCR Mix, 0.5 µL 40× RT Enzyme Mix, 1 µL each of 20× TaqMan Gene Expression Assays for the target gene (FAM) and GAPDH control (VIC), 6.5 µL RNase-free water, and 1 µL cfRNA. Assays used were CEACAM6 (Hs03645554_m1), MUC1 (Hs00159357_m1), SFTPB (Hs00167036_m1), SFTPA1 (Hs00831305_s1), SLC34A2 (Hs00197519_m1), MBP (Hs00921945_m1), and GAPDH (Hs99999905_m1). Thermal cycling was performed on a CFX96 Real-Time PCR Detection System (Bio-Rad) with reverse transcription at 48 °C for 15 min, polymerase activation at 95 °C for 10 min, followed by 40 cycles of 95 °C for 15 s and 60 °C for 1 min. Relative expression was calculated using the comparative Ct method following normalization to GAPDH. Technical replicates were averaged prior to analysis.

### Statistical Analyses

Statistical analyses were performed using R (v4.4.1). Principal component analysis and hierarchical clustering were performed using rlog-transformed counts. Unless otherwise specified, statistical tests were two-sided and significance was defined as an adjusted p value < 0.05 following Benjamini–Hochberg false discovery rate (FDR) correction. Nonparametric statistical tests were used for group comparisons, including Kruskal-Wallis and Wilcoxon rank-sum tests, due to modest sample sizes and heterogeneity in patient-derived data without assuming normal distributions. For analyses involving repeated sampling, matched samples, or technical replicates, appropriate statistical approaches were applied. Longitudinal CSF samples collected from the same patient at multiple time points were treated as repeated measures, with each sample representing a distinct biological specimen. Matched inter-site analyses were performed using paired statistical tests. Technical replicates for RT–qPCR were averaged prior to statistical testing, with each biological sample treated as the unit of analysis. No experimental randomization or investigator blinding was performed.

## Data Availability

Gene count matrices, associated metadata, and sequencing data will be deposited in GEO. All other data supporting the findings of this study are available within the Article, Supplementary Information, or Source Data.

## ACKNOWLEDGMENTS

The authors thank Stanford Health Center patients (and their families) for donating tissue for research purposes.

## AUTHOR CONTRIBUTIONS

M.U.G., G.B., and M.H.G. designed the study. M.H.G. supervised the study. M.U.G., G.B., T.T.M.N, and M.H.G. designed experiments. R.T., D.H., G.Br., C.R-C., G.C., E.V., M.G., T.B., and S.N. contributed to sample collection. M.U.G., G.B., P.N.P., T.T., R.T., D.H., M.R.O., S.L., C.W., G.Br., B.G., S.C., and S.T. performed experiments and data collection. M.U.G., G.B., P.N.P, C.W. performed data analysis. M.U.G., G.B., P.N.P, and Y.Z. wrote manuscript with input from all authors. All authors reviewed and approved the final manuscript.

