## Supplementary Figures 1 - 9 for "CSF-Seq enables transcriptome-wide profiling of cerebrospinal fluid and identifies prognostic signature of leptomeningeal disease"

### SUPPLEMENTARY FIGURE LEGENDS

#### Supplemental Figure 1

**a**

**Global RNA Sequencing Metrics (n = 129)**

| Description | Mean | Median | Min | Max |
|---|---|---|---|---|
| Input Reads | 118,360,801 | 112,176,381 | 41,985,721 | 287,858,599 |
| Uniquely Mapped Reads | 23,547,840 | 19,327,739 | 167,163 | 112,083,509 |
| Exon Fraction | 0.54 | 0.54 | 0.064 | 0.87 |
| Intron Fraction | 0.21 | 0.19 | 0.069 | 0.68 |
| Intergenic Fraction | 0.25 | 0.25 | 0.028 | 0.52 |

**b**

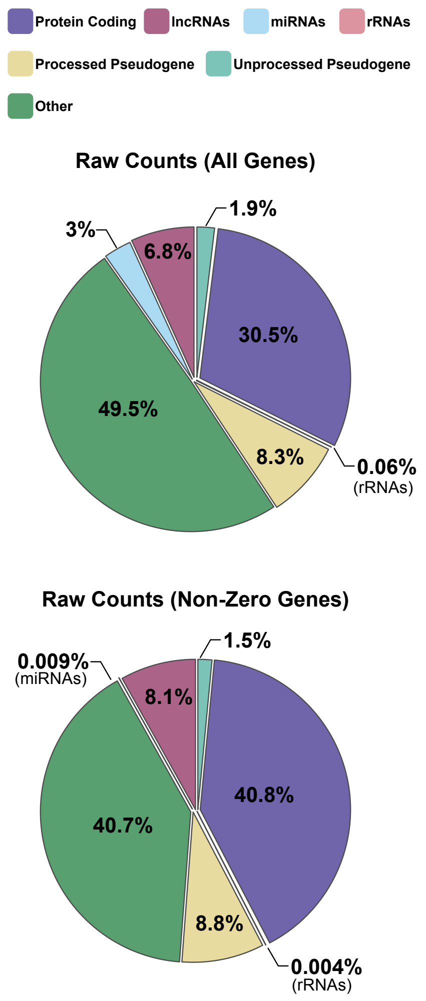


**c**

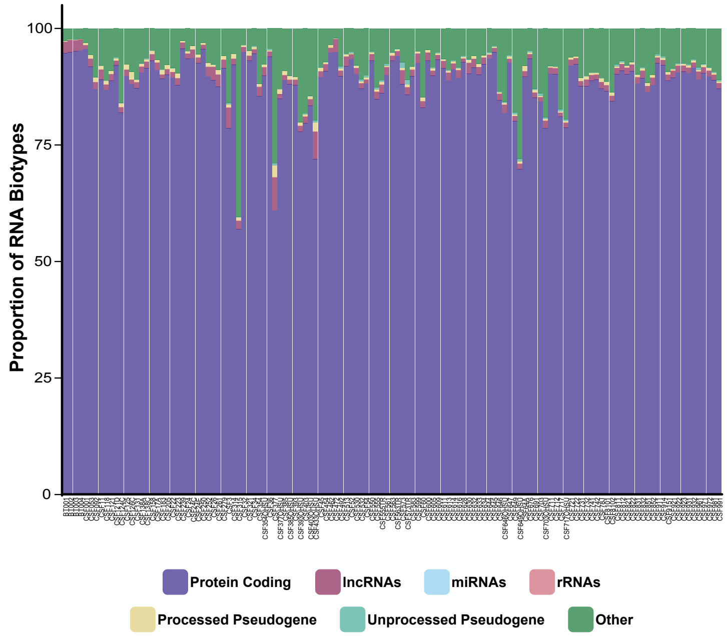


**d** **e** **f**

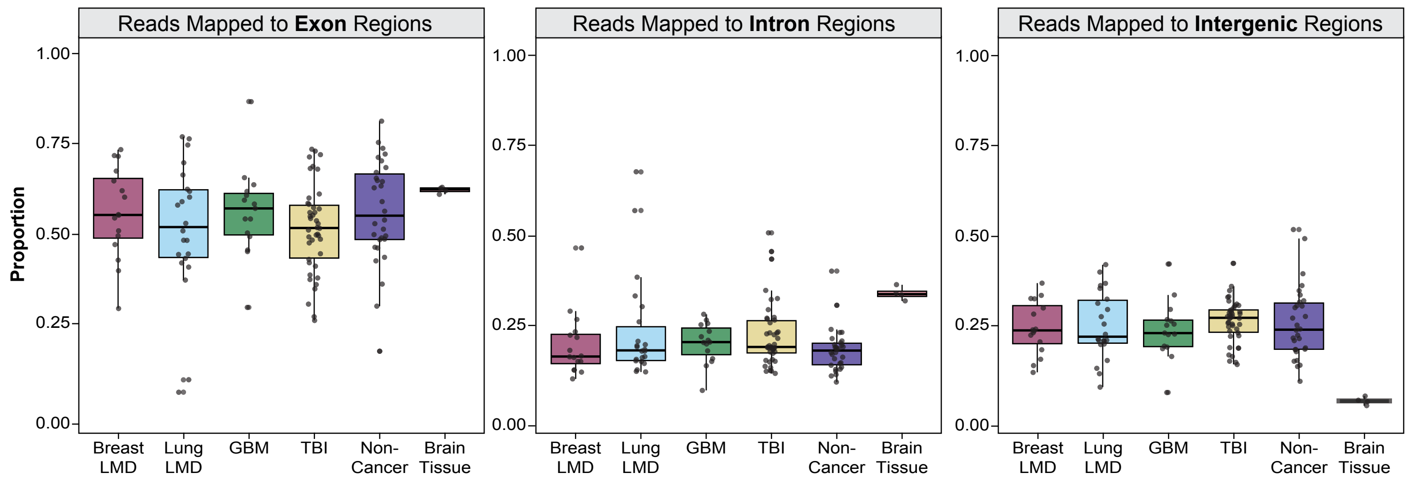


**Supplementary Figure 1. Global sequencing metrics and RNA composition across CSF-Seq samples. (a)** Summary table of global RNA sequencing metrics across the full cohort (n = 129 samples), including input reads, uniquely mapped reads, and the proportion of reads mapping to exonic, intronic, and intergenic regions. Mean, median, minimum, and maximum values are shown. **(b)** Distribution of RNA biotypes across all samples. Pie charts display the proportion of raw counts assigned to annotated RNA biotypes, shown for all detected genes (top) and non-zero genes (bottom). Categories include protein-coding, long non-coding RNA (lncRNA), microRNA (miRNA), ribosomal RNA (rRNA), processed pseudogene, unprocessed pseudogene, and other annotated transcripts. **(c)** Per-sample RNA biotype composition. Stacked bar plots display the proportion of reads assigned to each RNA biotype per sample (n = 129); each bar represents one CSF sample. **(d-f)** Distribution of reads mapped to genomic features across disease categories. Boxplots show the proportion of reads mapping to **(d)** exonic regions, **(e)** intronic regions, and **(f)** intergenic regions; each dot represents one CSF sample. No statistical tests were performed; plots are descriptive. For boxplots, the center line denotes the median, box edges represent the 25th and 75th percentiles, and whiskers extend to ±1.5× interquartile range (IQR).

### Supplemental Figure 2

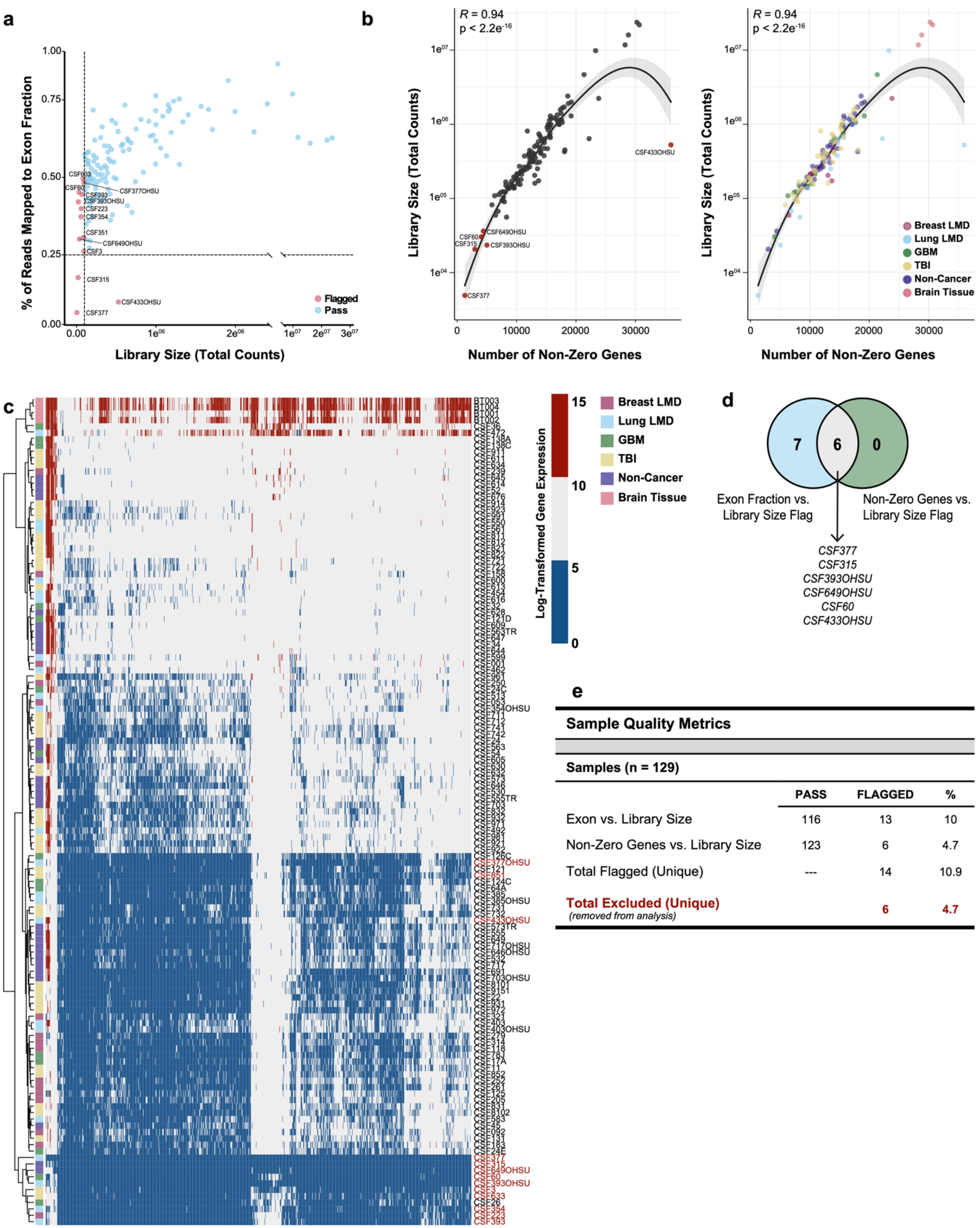


**Sample Quality Metrics**

**Samples (n = 129)**

| | PASS | FLAGGED | % |
|---|---|---|---|
| Exon vs. Library Size | 116 | 13 | 10 |
| Non-Zero Genes vs. Library Size | 123 | 6 | 4.7 |
| Total Flagged (Unique) | --- | 14 | 10.9 |
| **Total Excluded (Unique)** *(removed from analysis)* | | **6** | **4.7** |

**Supplementary Figure 2. Multi-parameter quality control (QC) and sample filtering prior to downstream transcriptomic analyses. (a)** Scatterplot of library size (total counts) versus exon fraction for all samples (n = 129); each dot represents one CSF sample. Samples flagged based on exon fraction versus library size criteria are shown in pink and labeled; samples passing QC are shown in blue. **(b)** Scatterplots of number of detected genes (non-zero genes) versus library size. Left: samples flagged based on non-zero gene detection criteria are highlighted in red and labeled. Right: samples colored by disease group. Coefficient of determination from linear regression ($R^2$) and associated two-sided p value are indicated. **(c)** Heatmap of log-transformed gene expression across samples (n = 129). Samples are hierarchically clustered based on global gene expression patterns; low-quality libraries are labeled in red. **(d)** Venn diagram summarizing overlap between orthogonal filtering criteria (exon fraction versus library size and non-zero genes versus. library size). **(e)** Summary table of QC metrics across the cohort. PASS and FLAGGED counts are shown for each filtering method. A total of 14 samples (10.9%) were flagged by at least one criterion; 6 unique samples (4.7%) were excluded from downstream analyses. **CSF**, cerebrospinal fluid; **GBM**, glioblastoma; **LMD**, leptomeningeal disease; **TBI**, traumatic brain injury.

### Supplemental Figure 3

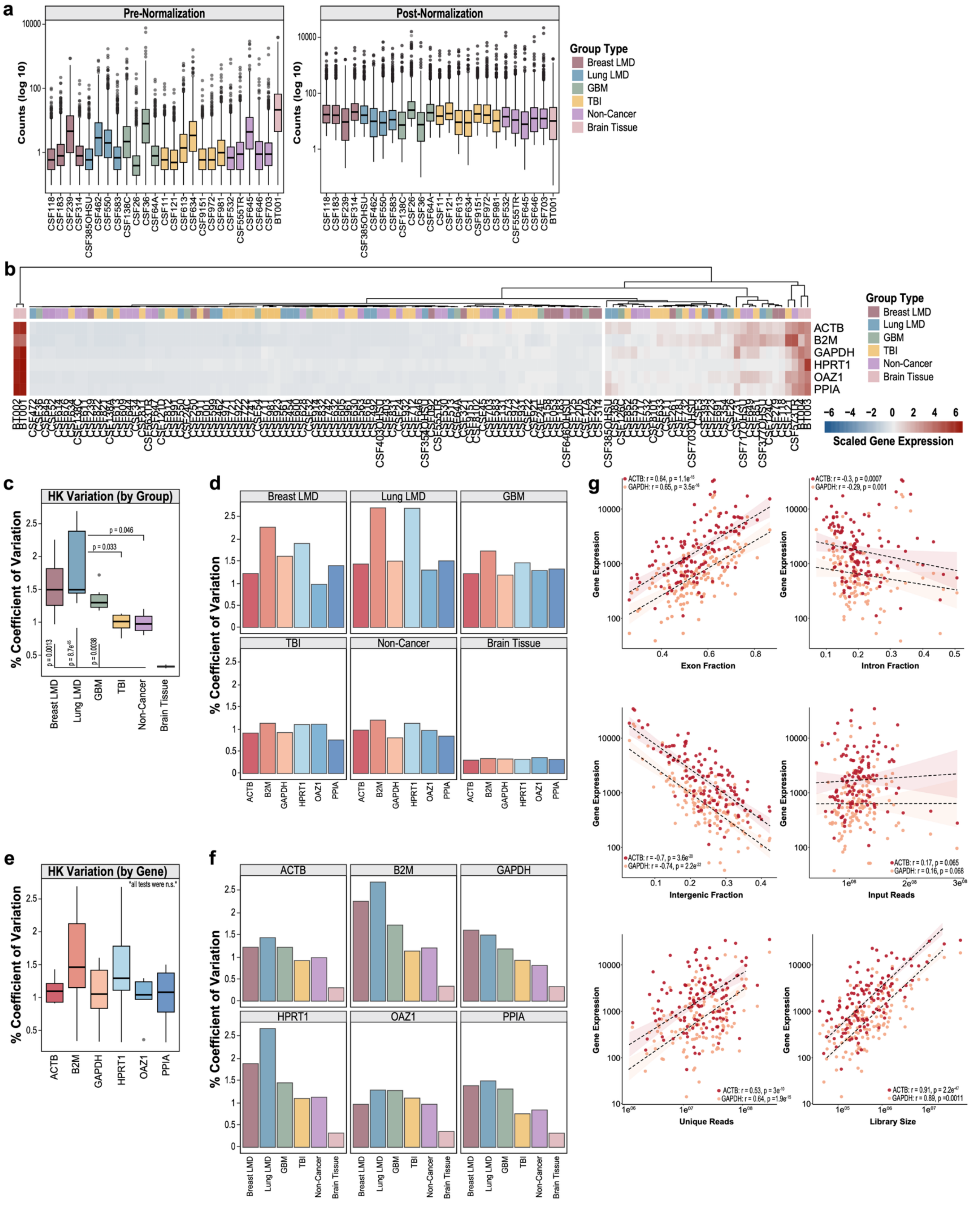


**Supplementary Figure 3. Normalization performance and evaluation of housekeeping gene stability across CSF samples. (a)** Global gene count distributions before and after normalization. Boxplots show log10 gene counts for all detected protein-coding genes (n = 18,649) across randomly selected samples from each disease group (breast

LMD, n = 4; lung LMD, n = 4; GBM, n = 4; TBI, n = 7; non-cancer, n = 5; brain tissue, n = 1); each dot represents one protein-coding gene. Left: pre-normalization counts. Right: DESeq2-normalized counts. **(b)** Heatmap of expression for six housekeeping genes (ACTB, B2M, GAPDH, HPRT1, OAZ1, PPIA) across all samples (n = 123). Gene expression values are scaled by row. Samples are hierarchically clustered and annotated by disease group. **(c)** Percent coefficient of variation (CV) of housekeeping genes within each disease group. For each group, CV values were calculated for the six housekeeping genes across samples in that group. Boxplots display gene-level CV distributions per disease group. **(d)** Percent CV of each housekeeping gene stratified by disease group. **(e)** Percent CV for each housekeeping gene calculated across all samples (n = 123). Boxplots display sample-level CV distributions for each gene. **(f)** Percent CV across disease groups stratified by housekeeping gene. **(g)** Correlation analyses between housekeeping gene expression (ACTB, red; GAPDH, orange) and sequencing metrics, including exon fraction, intron fraction, intergenic fraction, input reads, uniquely mapped reads, and library size; each dot represents one CSF sample (n = 123). Pearson correlation coefficient (r) and two-sided p value are shown. Linear regression lines with 95% confidence intervals are displayed. For **(c)** and **(e)**, p values were calculated using two-sided Kruskal–Wallis tests with Dunn correction for multiple comparisons. For boxplots in **(a)** and **(c–f)**, the center line denotes the median, box edges represent the 25th and 75th percentiles, and whiskers extend to ±1.5× interquartile range (IQR). **CV**, coefficient of variation; **GBM**, glioblastoma; **LMD**, leptomeningeal disease; **TBI**, traumatic brain injury.

### Supplemental Figure 4

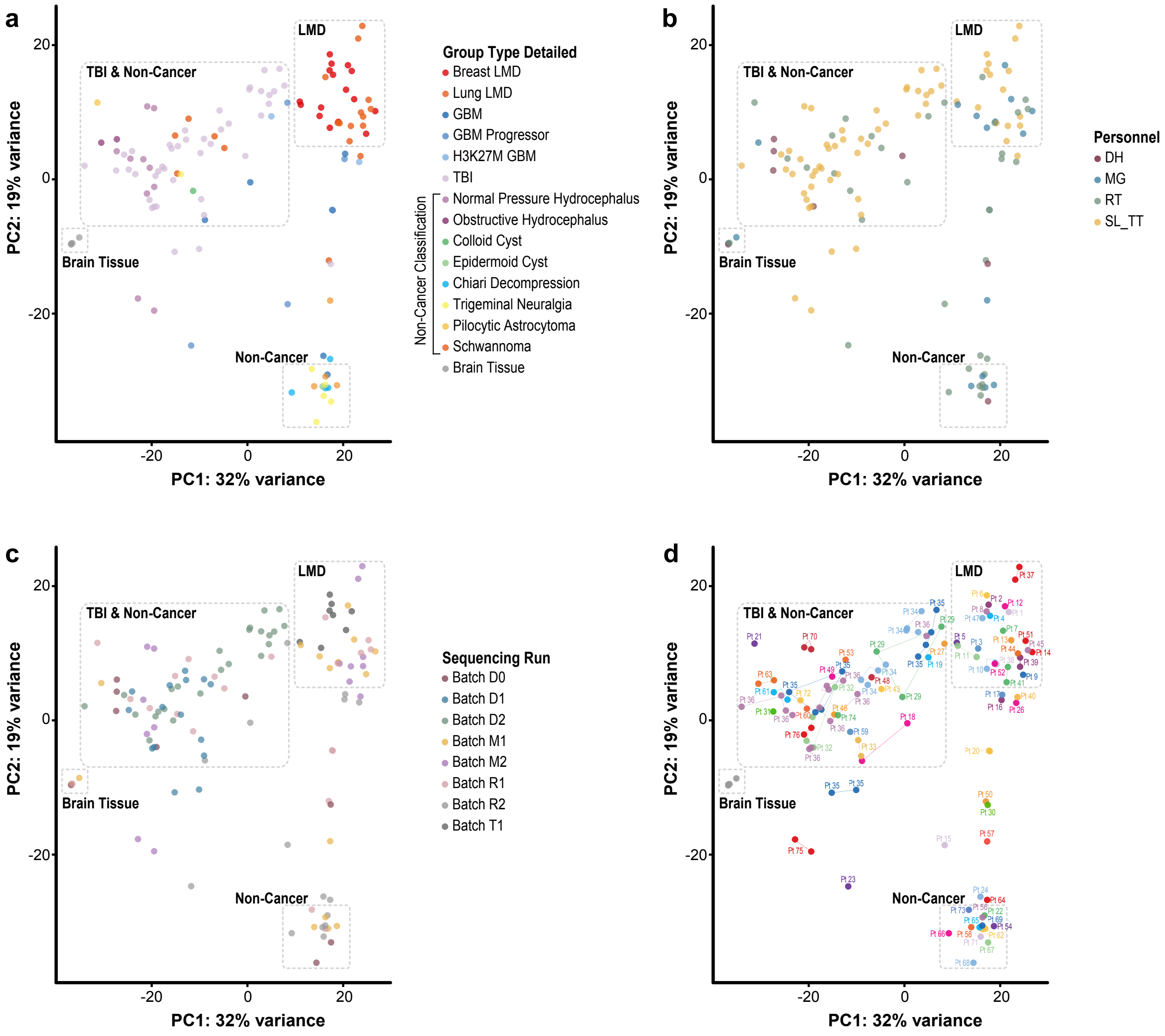


**Supplementary Figure 4. Principal component analyses annotated by clinical and technical variables. (a-d)** PCA of normalized protein-coding gene expression (n = 18,649 genes; n = 123 samples). **(a)** Samples annotated by detailed clinical diagnosis. **(b)** Samples annotated by processing personnel. **(c)** Samples annotated by sequencing batch. **(d)** Samples annotated by patient ID; samples from the same patient are connected by dotted lines. For plots **(a-d)**, each dot represents one CSF sample. Dotted outlines indicate disease groupings consistent with Fig. 3d–e. PC1 explains 32% of the variance and PC2 explains 19% of the variance. **PCA**, principal component analysis.

### Supplemental Figure 5

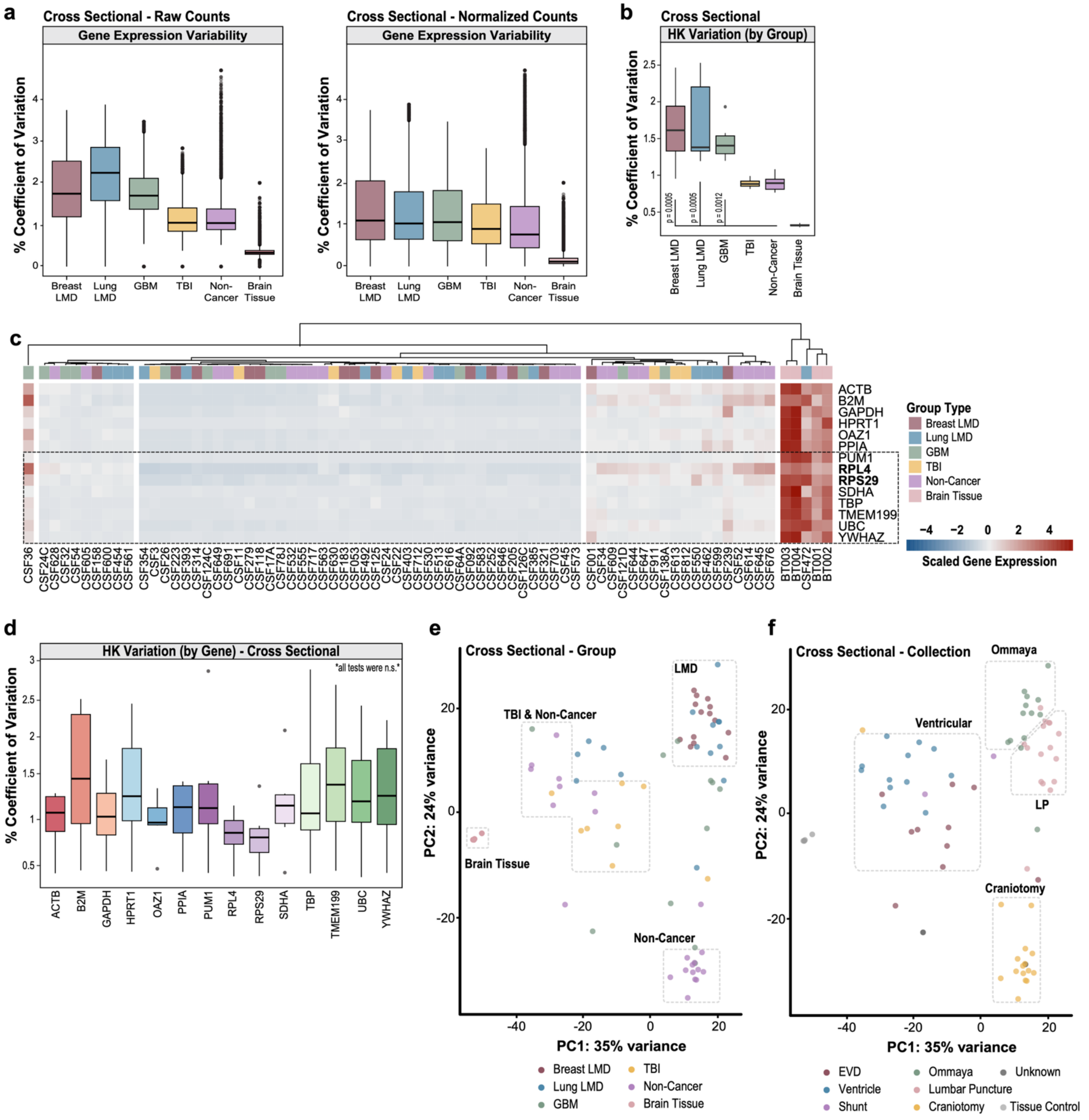


**Supplementary Figure 5. Cross-sectional analyses of transcriptome variability and housekeeping genes.** All analyses shown were performed using cross-sectional CSF samples only (n = 75; one CSF sample per patient). **(a)** Gene expression variability before and after normalization. Boxplots show the distribution of percent CV for protein-coding genes across disease groups using raw counts (left) and normalized counts (right); each dot represents one protein-coding gene. **(b)** Percent CV of six housekeeping genes (ACTB, B2M, GAPDH, HPRT1, OAZ1, PPIA) across disease groups in cross-sectional CSF samples. P values were calculated using two-sided Kruskal–Wallis with Dunn correction for multiple comparisons. **(c)** Heatmap of scaled gene expression for six canonical housekeeping genes (from panel b) and eight additional genes (PUM1, RPL4, RPS29, SDHA, TBP, TMEM199, UBC, YWHAZ). Gene expression values are scaled by row. Samples (n = 75) are hierarchically clustered and annotated by disease group. **(d)** Percent CV of fourteen candidate housekeeping genes calculated across all cross-sectional CSF samples (n = 75); each dot represents one CSF sample. **(e-f)** PCA of normalized protein-coding gene expression (n = 18,649 genes;

n = 75 samples); each dot represents one CSF sample. PC1 explains 35% of the variance and PC2 explains 24% of the variance. **(e)** Samples annotated by disease group. **(f)** Samples annotated by collection modality. For boxplots, the center line denotes the median, box edges represent the 25th and 75th percentiles, and whiskers extend to ±1.5× interquartile range (IQR). **CV**, coefficient of variation; **GBM**, glioblastoma; **HK**, housekeeping; **LMD**, leptomeningeal disease; **LP**, lumbar puncture; **PCA**, principal component analysis; **TBI**, traumatic brain injury.

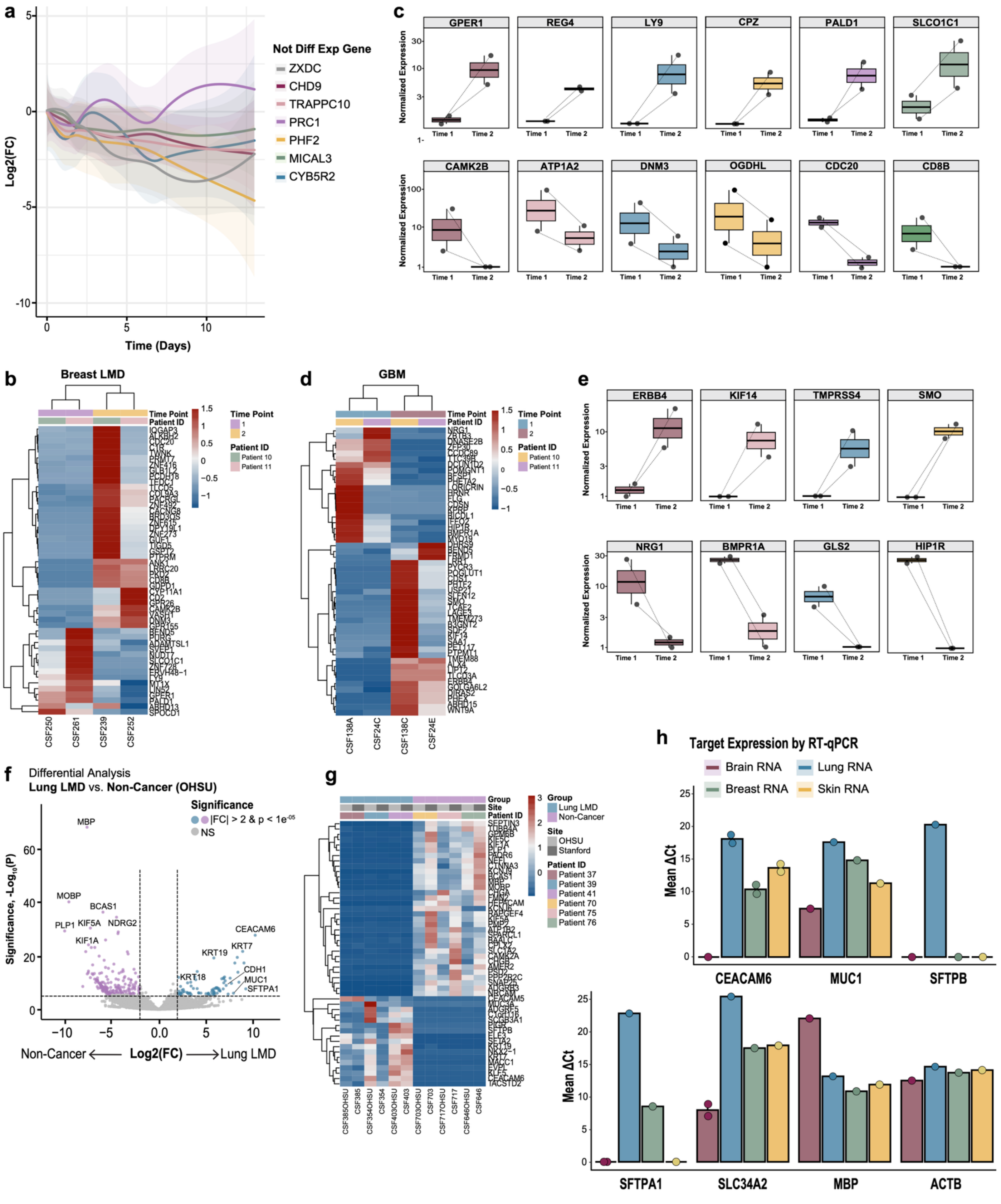

**Supplementary Figure 6. Longitudinal and inter-site analyses supporting temporal dynamics and reproducibility of CSF-Seq. (a)** Line plot shows representative non-differentially expressed transcripts across time (days 1, 2, and 10); curves represent LOESS-smoothed log2 fold change trajectories. Longitudinal CSF samples collected from the same patients at multiple time points were treated as repeated measures, with each time point representing a distinct biological sample. **(b)** Heatmap of longitudinal gene expression in matched breast LMD samples. Scaled normalized expression values are shown; genes are clustered by rows, and samples by columns. Samples are annotated by patient ID and time point. **(c)** Gene-level expression across time in matched breast LMD samples (n = 4). Boxplots show normalized expression values for selected genes across two time points; each dot represents one CSF sample. **(d)** Heatmap of longitudinal gene expression in matched GBM samples. Scaled normalized expression values are shown; genes are clustered by rows, and samples by columns. Samples are annotated by patient ID and time point. **(e)** Gene-level expression across time in matched GBM samples (n = 4). Boxplots show normalized expression values for selected genes across two time points; each dot represents one CSF sample. **(f)** Differential expression analysis comparing lung LMD versus non-cancer samples processed at OHSU using DESeq2. Volcano plot shows log2 fold change versus –log10(adjusted p value); each dot represents one protein-coding gene. Genes exceeding |log2 fold change| > 2 and adjusted $p < 1\times10^{-5}$ are colored. **(g)** Heatmap of scaled normalized expression across matched inter-site samples. Genes are clustered by rows. Samples are annotated by patient ID, processing site, and disease group. **(h)** Target expression measured by RT–qPCR across tissue RNA controls (brain, lung, breast, and skin RNA). Bar plots show mean ΔCt values; each dot represents one biological sample (averaged across technical replicates, n = 2-3). For boxplots, the center line denotes the median, box edges represent the 25th and 75th percentiles, and whiskers extend to ±1.5× interquartile range (IQR). **GBM**, glioblastoma; **LMD**, leptomeningeal disease; **LOESS**, locally estimated scatterplot smoothing; **RT–qPCR**, reverse transcription quantitative PCR; **TBI**, traumatic brain injury.

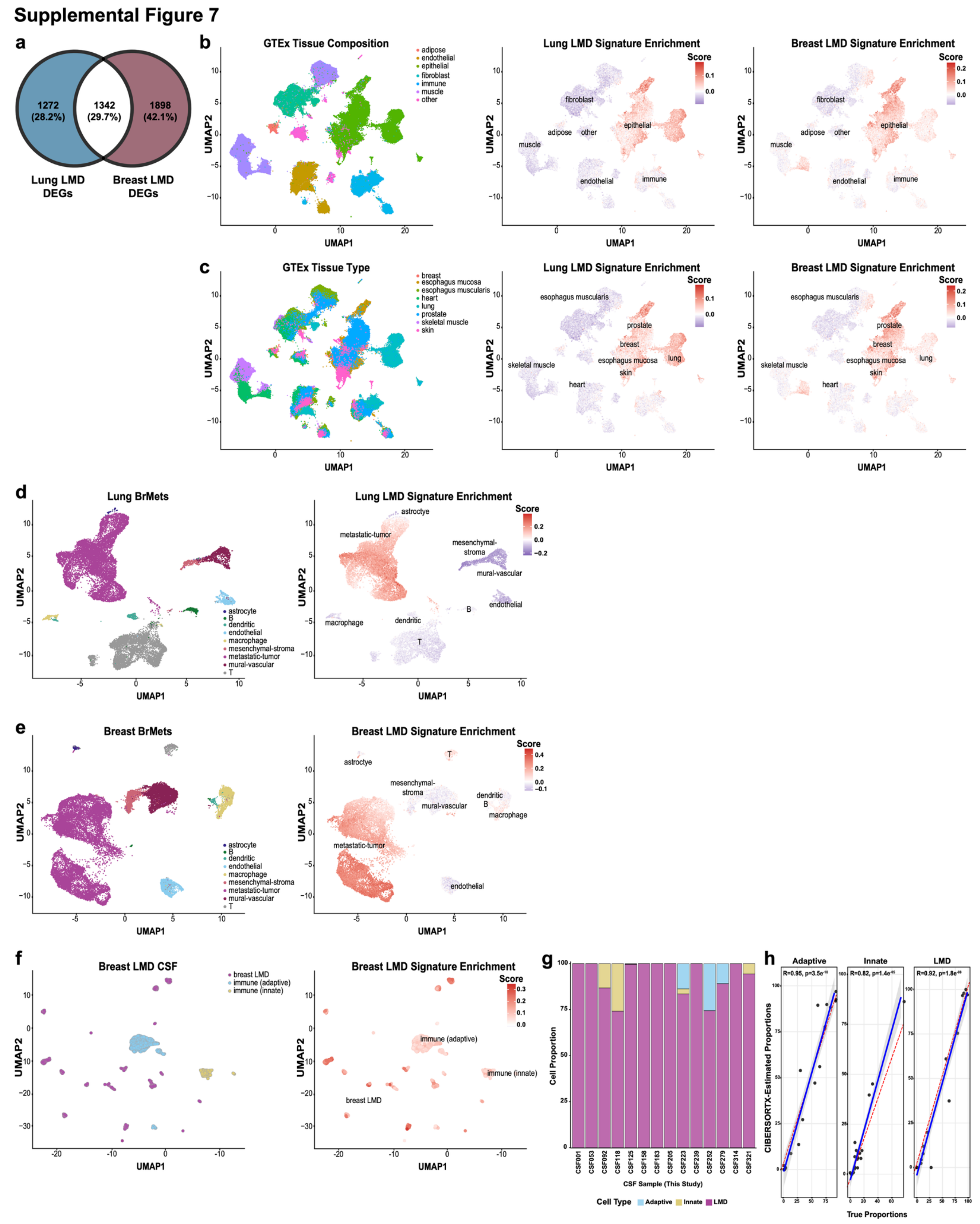


**Supplementary Figure 7. External validation and cellular context of LMD-associated transcriptional programs. (a)** Venn diagram showing overlap of differentially expressed genes between breast LMD (versus non-cancer), and lung LMD (versus non-cancer). **(b)** Projection of LMD signature scores onto GTEx tissue composition single-cell reference UMAP. Left: cells annotated by tissue composition. Middle: cells annotated by lung LMD signature score. Right: cells annotated by breast LMD signature score. **(c)** Projection of LMD signature scores onto GTEx tissue type single-cell reference UMAP. Left: cells annotated by tissue type. Middle: cells annotated by lung LMD signature score. Right: cells annotated by breast LMD signature score. **(d)** Projection of lung LMD signature score onto lung brain metastasis single-cell dataset. Left: cells annotated by cell type. Right: cells annotated by lung LMD signature score. **(e)** Projection of breast LMD signature score onto breast cancer brain metastasis single-cell dataset. Left: cells annotated by cell type. Right: cells annotated by breast LMD signature score. **(f)** Projection of breast LMD signature score onto breast LMD CSF single-cell dataset. Left: cells annotated by cell type. Right: cells annotated by breast LMD signature score. **(g)** Deconvolution of breast LMD bulk CSF-Seq samples using single-cell reference shown in (f). Bar plots display estimated cell-type proportions per sample; each bar represents one CSF sample. **(h)** Correlation between known cell-type proportions and CIBERSORTx-estimated proportions for adaptive immune, innate immune, and breast LMD cell populations; each dot represents one CSF sample. Pearson correlation coefficient (r) and associated two-sided p value are shown. For **(b-f)**, each dot represents one single cell. **GTEx**, Genotype-Tissue Expression; **LMD**, leptomeningeal disease; **UMAP**, Uniform Manifold Approximation and Projection.

### Supplemental Figure 8

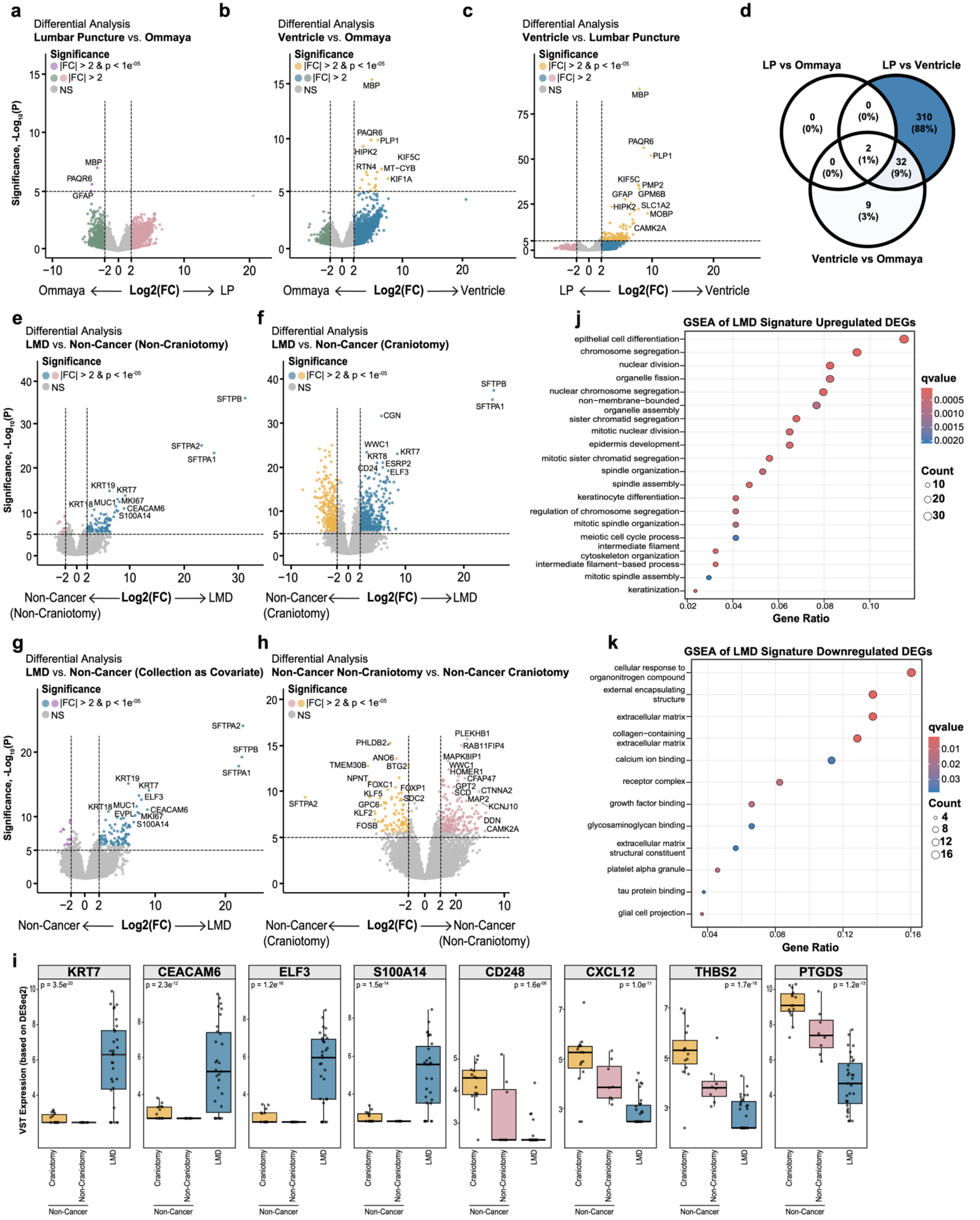


**Supplementary Figure 8. Evaluation of collection modality effects and multistep derivation of collection-**

**independent LMD signature. (a-c)** Differential expression analyses comparing lung LMD CSF samples by collection modality using DESeq2: **(a)** lumbar puncture (LP; n = 9) versus Ommaya (n = 2), **(b)** ventricle (n = 6) versus Ommaya (n = 2), and **(c)** ventricle (n = 6) versus LP (n = 9). Volcano plots show log2 fold change versus –log10(adjusted p value); each dot represents one protein-coding gene. Genes exceeding |log2 fold change| > 2 and adjusted $p < 1\times10^{-5}$ are colored; NS, not significant. **(d)** Venn diagram summarizing overlap of differentially expressed genes across collection-modality comparisons from (a-c). **(e-g)** Differential expression analyses comparing LMD versus non-cancer controls under different modeling strategies using DESeq2: **(e)** LMD (n = 31) versus non-cancer non-craniotomy samples only (n = 8), **(f)** LMD (n = 31) versus non-cancer craniotomy samples only (n = 15), and **(g)** LMD (n = 31) versus full cohort of non-cancer (n = 23), including collection modality as covariate in DESeq2 model. Volcano plot shows log2 fold change versus –log10(adjusted p value); each dot represents one protein-coding gene. Genes exceeding |log2 fold change| > 2 and adjusted $p < 1\times10^{-5}$ are colored; NS, not significant. **(h)** Differential expression analysis comparing non-cancer craniotomy (n = 15) versus non-cancer non-craniotomy (n = 8) samples using DESeq2. Volcano plot shows log2 fold change versus –log10(adjusted p value); each dot represents one protein-coding gene. Genes exceeding |log2 fold change| > 2 and adjusted $p < 1\times10^{-5}$ are colored; NS, not significant. **(i)** Expression of representative genes retained in the final LMD signature across LMD and non-cancer samples stratified by collection modality. Boxplots show normalized expression values; each dot represents one CSF sample. For boxplots, the center line denotes the median, box edges represent the 25th and 75th percentiles, and whiskers extend to ±1.5× interquartile range (IQR). (**j-k**) GSEA of **(j)** genes upregulated and **(k)** genes downregulated in the final 460-gene LMD signature. NES and adjusted p values (q values) are shown. **DEG**, differentially expressed genes; **GSEA**, gene set enrichment analysis; **LMD**, leptomeningeal disease; **LP**, lumbar puncture; **NES**, normalized enrichment score.

### Supplemental Figure 9

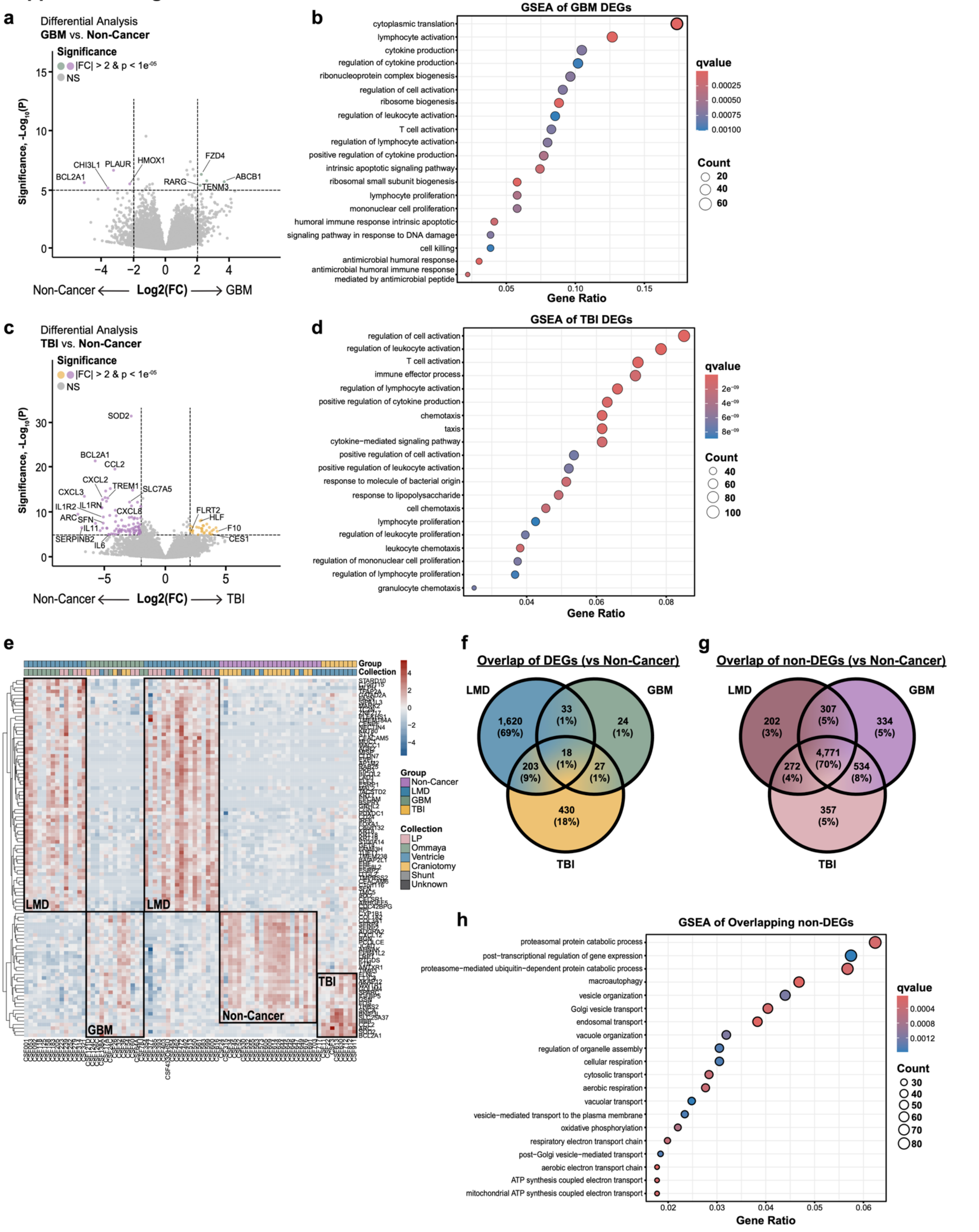


**Supplementary Figure 9. Disease-specific transcriptional programs across neurological and oncologic CSF**

**samples. (a)** Differential expression analysis comparing GBM (n = 13) versus non-cancer CSF (n = 23) using DESeq2. Volcano plot shows log2 fold change versus –log10(adjusted p value); each dot represents one protein-coding gene. Genes exceeding |log2 fold change| > 2 and adjusted $p < 1\times10^{-5}$ are colored; NS, not significant. **(b)** GSEA of differentially expressed genes identified in (a). NES and adjusted p values (q values) are shown. **(c)** Differential expression analysis comparing TBI (n = 8) versus non-cancer CSF (n = 23) using DESeq2. Volcano plot shows log2 fold change versus –log10(adjusted p value); each dot represents one protein-coding gene. Genes exceeding |log2 fold change| > 2 and adjusted $p < 1\times10^{-5}$ are colored. **(d)** GSEA of differentially expressed genes identified in (c). NES and adjusted p values (q values) are shown. **(e)** Heatmap of normalized expression values across all disease groups. Genes are clustered by rows. Samples are annotated by disease category. Highlighted regions indicate disease-specific transcriptional clusters. **(f)** Venn diagram showing overlap of differentially expressed genes across LMD, GBM, and TBI comparisons. **(g)** Venn diagram showing overlap of non-differentially expressed genes across LMD, GBM, and TBI comparisons. **(h)** GSEA of non-differentially expressed genes across disease groups from (g). NES and adjusted p values (q values) are shown. **DEG**, differentially expressed genes; **GBM**, glioblastoma; **GSEA**, gene set enrichment analysis; **LMD**, leptomeningeal disease; **NES**, normalized enrichment score; **TBI**, traumatic brain injury.
